# Disruption of ADNP-KDM1A-GTF2I complex drives neural differentiation imbalance in Helsmoortel-Van der Aa syndrome

**DOI:** 10.1101/2025.03.06.641037

**Authors:** Ludovico Rizzuti, Alessandro Vitriolo, Pelicano de Almeida Mariana, Marlene F. Pereira, Lize Meert, Francesco Dossena, Filippo Prazzoli, Mike Dekker, Dick H.W. Dekkers, Jeroen Demmers, Wilfred F.J. van IJcken, Chiara Soriani, Michele Gabriele, Mohiuddin Mohiuddin, Anke Van Dijck, Veronica Finazzi, Sebastiano Trattaro, Danila Pallotta, Erika Tenderini, Anneke T. Vulto-van Silfhout, Bert B. A. de Vries, Evan E. Eichler, Christopher E. Pearson, R. Frank Kooy, Raymond Poot, Giuseppe Testa

## Abstract

Mutations in ADNP (Activity-Dependent Neuroprotective Protein) are among the most frequent monogenic causes of autism spectrum disorder (ASD) and lead to Helsmoortel-Van der Aa syndrome (HVDAS). Yet how ADNP dysfunction leads to HVDAS is unclear. We employed patient-derived induced pluripotent stem cells, cortical organoids and ADNP KO human neural stem cells (hNSCs) to clarify the cellular and molecular mechanism of HVDAS onset. We purified an ADNP-KDM1A-GTF2I (AKG) protein complex from hNSCs and show that it targets transposable elements (TEs) to repress nearby gene transcription. Upon ADNP KO, KDM1A binding is lost at promoters targeted by AKG, pointing to ADNP as the anchoring subunit of the AKG complex. HVDAS cortical organoids show impaired progenitor proliferation and accelerated neuronal differentiation, coupled with a sustained upregulation of neurogenesis transcriptional programs, including key transcription factors normally repressed by AKG. This work suggests that the AKG complex acts as the relevant ADNP unit in the molecular onset of HVDAS.

## INTRODUCTION

Neurodevelopmental disorders (NDD) typically manifest during early childhood and are characterised by a varying degree of neuropsychiatric impairments, often coupled with multi-organ manifestations^1,2^. The study of NDDs has benefited from the detection and characterisation of associated genomic variants, enabling an improved resolution in establishing causative links between genotype and phenotype^3,4^. Rare inherited and *de novo* mutations in chromatin and transcriptional regulators can cause autism spectrum disorder (ASD)^5–9^. Among genes most frequently *de novo* mutated in ASD is Activity Dependent Neuroprotective Protein (ADNP)^10–13^. ADNP mutations cause Helsmoortel-Van der Aa syndrome (HVDAS), an autosomal-dominant NDD featuring intellectual disability along with ASD, developmental delay, verbal impairment, and facial dysmorphisms^14–16^. ADNP encodes for a transcription factor (TF) protein originally found to cause neural tube closure defects when deficient in mouse embryos^17,18^. Molecular characterisation of ADNP in mouse embryonic stem cells (mESC) showed that it constitutes the DNA-binding subunit of the ChAHP complex, which also includes CHD4 and HP1γ^19^. ChAHP localises to euchromatic regions where it can establish inaccessible chromatin domains independently of H3K9me3. This mode of repression prevents premature gene activation during neuroectodermal specification^19^. Moreover, the ChAHP complex was reported to compete with CTCF for a subset of common binding motifs, suggesting a role for ADNP in shaping mammalian genome organization^20^. ADNP was also shown to recruit TFIIIC to create contacts between distal genes to maintain their poised state for reactivation^21^.

Here, we established an integrated resource including: i) CRISPR-Cas9-edited induced pluripotent stem cells (iPSCs) and neural stem cells (hNSCs) models, featuring endogenously tagged and knock-out ADNP alleles; ii) iPSC-derived cortical brain organoid models from a uniquely informative cohort of ADNP patients with representative mutations. We found that ADNP predominantly binds to cis-regulatory elements in TEs to mostly repress gene expression, and that its absence correlates with increased local levels of activating histone marks. We purified ADNP complexes from hNSCs and identified through mass spectrometry a novel complex including ASD-related chromatin factors KDM1A and GTF2I, that we named AKG complex. Finally we observe that HVDAS cortical organoids show altered proliferative behaviour and aberrant cell type abundance, pointing to an accelerated neuronal differentiation. We link this molecular HVDAS phenotype with the activity of AKG, which we find to repress neurogenesis-related transcriptional networks in cortical brain organoids.

## RESULTS

### ADNP dosage impairment alters early developmental lineages

To understand how ADNP mutations affect transcriptional regulation and cell fate decisions, we generated iPSC lines from six HVDAS patients (three males and three females) carrying *de novo* pathological frameshift or nonsense mutations spanning the ADNP gene, including the most common ADNP variant p.Tyr719*^14,22^ (**Fig.1A**, **Table S1**). Transcriptional profiling confirmed the canonical expression of pluripotency hallmarks (POU5F1, SOX2, NANOG, LIN28) (**Fig.S1A**). We quantified the expression levels of ADNP transcript using allele-specific digital PCR and observed equivalent expression of both *ADNP* alleles in all iPSC lines (**Fig.S1B**), showing that mutant alleles do not undergo appreciable nonsense mediated mRNA decay. We also generated ADNP-KO human neural stem cells (hNSCs) using CRISPR/Cas9 to excise approximately 1,800 base pairs (bp) of the coding portion of the ADNP gene, in homozygosity (**Fig.1A** and **Fig.S1C,D**). The absence of ADNP protein in two independently generated ADNP mutant lines was confirmed by Western Blot (**Fig.S1E**). Bulk RNA-seq of iPSC lines revealed 440 differentially expressed genes (DEGs) (FDR<0.05, fold change (FC)>1.5) between HVDAS lines and controls, with HVDAS iPSCs showing a prevalent upregulation (307 upregulated genes and 133 downregulated). RNA-seq performed on ADNP-KO hNSCs showed 458 upregulated genes and 373 downregulated genes compared to WT hNSCs (FDR<0.05, FC>2, **Table S2**). The majority of DEGs shared by iPSCs and hNSCs were upregulated (69,32% and 54,62% of DEGs, respectively), and their intersection (56 genes with FC 1.5) was significant (hypergeometric test, p=5.36e-05) (**Fig.1B**), uncovering a common subset of targets between the two cell types, most of which are derepressed upon ADNP loss. Compared to controls, HVDAS iPSCs showed a global increase in the average signal of H3K27ac peaks but not of H3K4me1 peaks (**Fig.1C, Fig.S1F**). Enhancers of upregulated DEGs in HVDAS iPSCs gained instead both H3K27ac and H3K4me1 signal (**Fig.1D,E**) more than downregulated DEGs (**Fig.S1F, S1G**) (H3K27ac gain p=4.41e-05, H3K4me1 gain p=6.19e-07, hypergeometric test), indicating that ADNP mutations result in the activation of specific cis-regulatory elements. Gene ontology (GO) analysis of DEGs in iPSCs revealed a high enrichment of genes related to differentiation and morphogenesis, while DEGs in hNSCs mostly enriched for basic cell functions such as cell-cell adhesion and DNA replication (**Fig.1F**). To probe how ADNP controls gene expression in these cell types, we annotated DEGs to identify known TFs that co-regulate ADNP targets. Master regulator analysis showed that the majority of DEGs in iPSCs and hNSCs are shared targets of other transcription factors, including KDM1A and GTF2I, suggesting that ADNP is part of a broader transcriptional regulatory network at both the pluripotent and neural stem cell states (**Fig.1H,I**). HVDAS patients often present craniofacial abnormalities, thus we also investigated the transcriptome of iPSC-derived neural crest stem cells (NCSCs). This cell population gives rise to peripheral neurons, chondrocytes, melanocytes and cells composing the craniofacial scaffold. We employed an established protocol to generate homogenous NCSCs cultures from four HVDAS lines and three control iPSC lines^23^. We assessed the acquisition of neural crest morphology through fluorescence-activated cell sorting (FACS) for known NCSC markers, confirming pure p75+ (NGFR) and HNK1+ populations after 21 days of differentiation (**Fig.S1H**). Bulk RNA-seq detected 149 upregulated and 105 downregulated (FDR < 0.05, FC>1.5) genes in HVDAS NCSCs compared to controls (**Fig.S1I**). GO analysis showed enrichment for categories such as axon development, synapse organisation and regulation of nervous system development, with the top enriched categories mostly including upregulated genes (**Fig.S1J**). Expectedly, the overlap between DEGs across all cell types (iPSC, NCS and NSC) was limited but interestingly their deregulation was largely explained by the same TFs (**Fig.S1K,L**), pointing to a convergent hub of ADNP dosage-sensitive genes whose dysregulation has a cell type-specific impact on gene expression programs.

**Figure 1.**
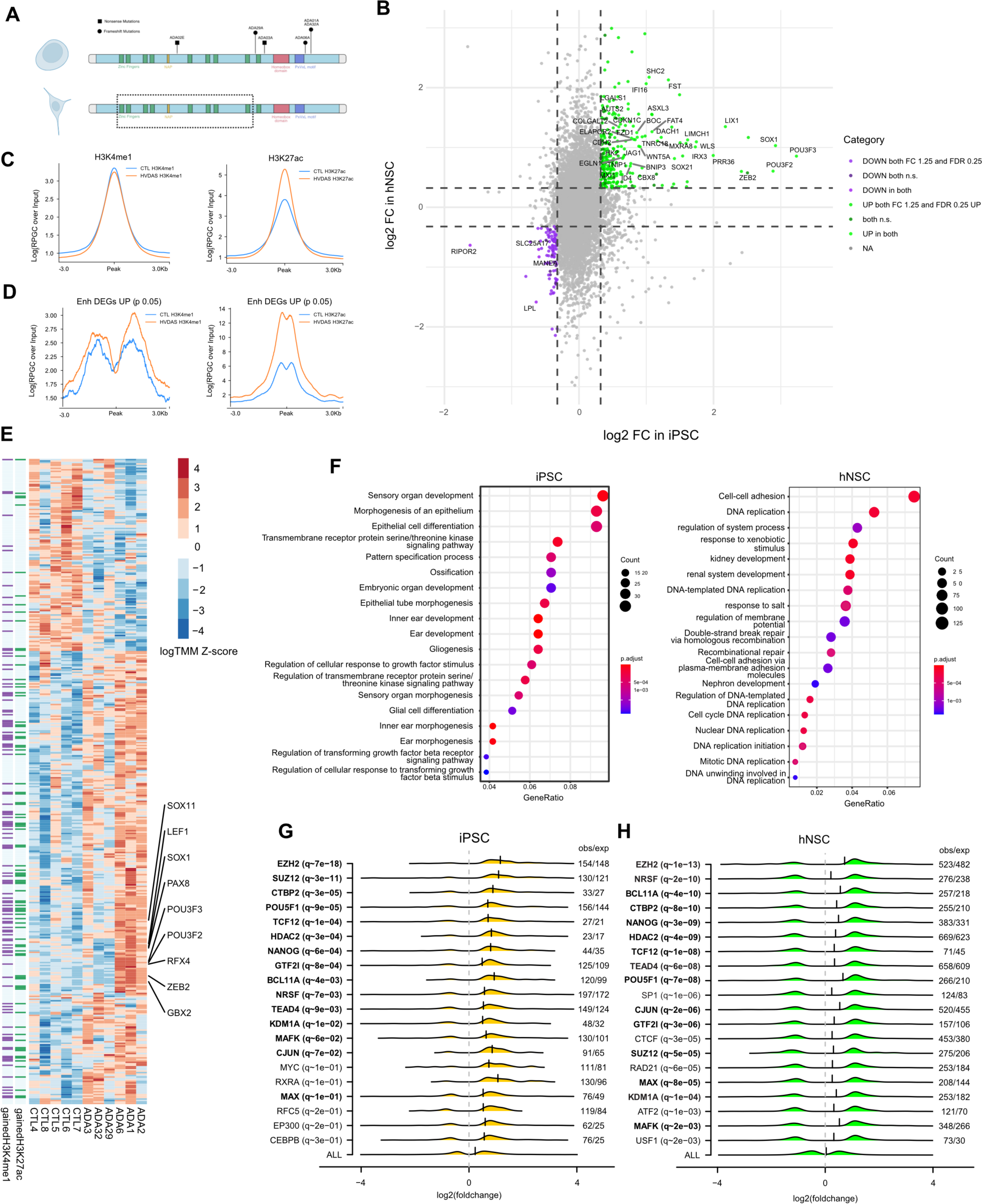
Gene dysregulation in HVDAS iPSCs and ADNP-KO hNSCs. **A.** Schematic representation of the protein domains of ADNP. For iPSCs (above) is reported the distribution of patients’ mutations, for NSC (below) is reported the excised portion in the ADNP-KO line (dashed rectangle). **B.** Scatter plot of differentially expressed genes in HVDAS iPSCs and ADNP KO hNSCs. log2(fold-change) measured in iPSC and hNSCs is respectively reported on the x-axis and y-axis. Purple color represents genes downregulated in both cell types. Green dots represent genes upregulated in both cell types. “DOWN in both” and “UP in both” represent genes significantly differentially expressed (FDR < 0.05 and FC > 1.5 in both cell types). Dots for genes with coherent fold change but limited significance (FDR < 0.25) or not significant (n.s., FDR >= 0.25) are reported with darker colors. To aid visualization only a portion of non-overlapping labels is reported for genes with FDR < 0.05 in both cell types (see the full list in **Table S2**). **C.** Global H3K27ac and H3K4me1 signal in HVDAS iPSCs (orange) and control iPSCs (blue). **D.** H3K27ac and H3K4me1 signal centered on enhancers of upregulated genes (p <0.05) in HVDAS iPSCs (orange) versus control iPSCs (blue). **E.** Heatmap of genes differentially expressed in iPSCs, genes that gained H3K27ac or H3K4me1 signal at regulatory regions are annotated on the left. Transcription factors that were upregulated and gained both marks are highlighted on the right. **F.** GO terms for RNA-seq in iPSCs (left) and RNA-seq in hNSCs (right); ‘Gene ratio’ indicates the portion of total DEGs in the given GO term. Dot size (Count) indicates the number of DEGs in each category. Dot color (p.adjust) represents significance of the GO categories. **G.** Master regulator analysis in iPSC shows the top transcription factors (TFs) based on gene enrichment analysis, by comparing DEGs with known sets of TF targets in pluripotent lineages; TFs are ordered by FDR (top to bottom); TFs in bold are significant in the master regulator analyses of both iPSC and hNSC. Along the x-axis we reported the log2(fold-change) distribution of DEGs targeted by each TF. **H.** Master regulator analysis in hNSCs. TFs are ordered by FDR (top to bottom); TFs in bold are significant also in the master regulator analysis of iPSC. Along the x-axis we reported the log2(fold-change) distribution of DEGs targeted by each TF.

### ADNP predominantly binds transposable elements to repress gene expression

We then investigated genome-binding by ADNP in both iPSCs and hNSCs. To overcome the lack of commercially available ChIP-grade antibodies for ADNP, we used CRISPR/Cas9 to endogenously incorporate, at the N-terminus of the ADNP open reading frame, a triple FLAG affinity tag (in iPSC) and a double FLAG and V5 affinity tag (in hNSCs) (Fig.2A). One heterozygous FLAG-tagged control iPSC line (hereafter referred to as CTL-FLAG), one homozygous FLAG-tagged patient iPSC line (carrying the mutation p.Tyr719*, hereafter referred to as HVDAS-FLAG) and one homozygous F2V5-ADNP hNSC control line were selected for further characterization (**Fig.S2A-D, Fig.S2E,F**). We detected the correctly sized protein in all lines but we did not find any evidence of the truncated form of the protein^24^. While CTL-FLAG ChIP-seq revealed ∼80,000 significant ADNP peaks (q<0.05), HVDAS-FLAG ChIP-seq showed a reduction in signal across all peaks (**Fig.2B**) and revealed ∼27,000 significant peaks, which almost completely overlapped with CTL-FLAG. In hNSCs, ADNP was present at ∼18,000 genomic sites, half of which (∼9,000 sites) were shared with CTL-FLAG (**Fig.2C, Fig.S2G**). The identified ADNP DNA-binding motif in iPSC lines and hNSCs revealed a shared consensus sequence that differs considerably from the CTCF motif previously reported in mESCs^20^ (**Fig.2D**). We performed ChIP-seq for CTCF in CTL-FLAG and HVDAS-FLAG iPSC lines (**Fig.S2H**), and found a comparable number of CTCF binding sites in both lines but only limited overlap with ADNP-bound regions (**Fig.S2J,K**), highlighting a fundamental difference between mouse and human ADNP biology. Coherently, ADNP KO and F2V5-ADNP hNSCs showed a limited overlap between ADNP and CTCF binding regions, and very limited changes in the binding distribution of CTCF (**Fig.S2I,L,M**). In agreement with previous results^19^, we observed ∼25% of ADNP sites positioned at promoters of protein-coding genes, while the majority of ADNP peaks (around 70%), in both iPSCs and hNSCs, fell in intronic and distal intergenic regions (**Fig.2E**). Interestingly, most ADNP sites overlapped with transposable elements (TEs) (**Fig.2F**), especially Alu sequences in iPSCs and MIR sequences in hNSCs (**Fig.2G**). Next, we investigated whether ADNP binding sites could account for the transcriptional dysregulation observed in HDVAS iPSCs and ADNP-KO hNSCs. First, we defined a set of iPSC regulatory elements by combining promoters of expressed genes with distal regions marked by both H3K4me1 and H3K27ac. For hNSC, in the absence of in-house generated H3K27ac and H3K4me1 we resorted to Roadmap Epigenomics, and used the same criteria used for hiPSC (see Methods). Among the 440 DEGs in iPSCs, ∼40% have a regulatory element targeted by ADNP (p=2.62e-26), with ADNP mostly targeting upregulated DEGs (∼80%, p=3.69e-230), consistent with a largely repressive role. Similarly, in hNSCs, ∼27% of DEGs have a regulatory element bound by ADNP (p=1.62e-40), of which 57% (p=4.05e-22) are upregulated (**Fig.2H**). Moreover, in both cell types, ADNP-bound regulatory regions of upregulated genes are enriched in TEs, both at promoters and enhancers (**Fig.2I**). Thus ADNP occupies regulatory elements of the majority of upregulated DEGs, which lose ADNP binding in HDVAS iPSCs, indicating that many DEGs are directly repressed by ADNP. Of note, ATAC-seq performed on HVDAS and control iPSCs did not reveal a significant difference in chromatin accessibility neither genome-wide, nor at TSS or at ADNP-lost sites (**Fig.S2N,O**) indicating that ADNP does not repress its target genes by creating local inaccessible chromatin domains.

**Figure 2.**
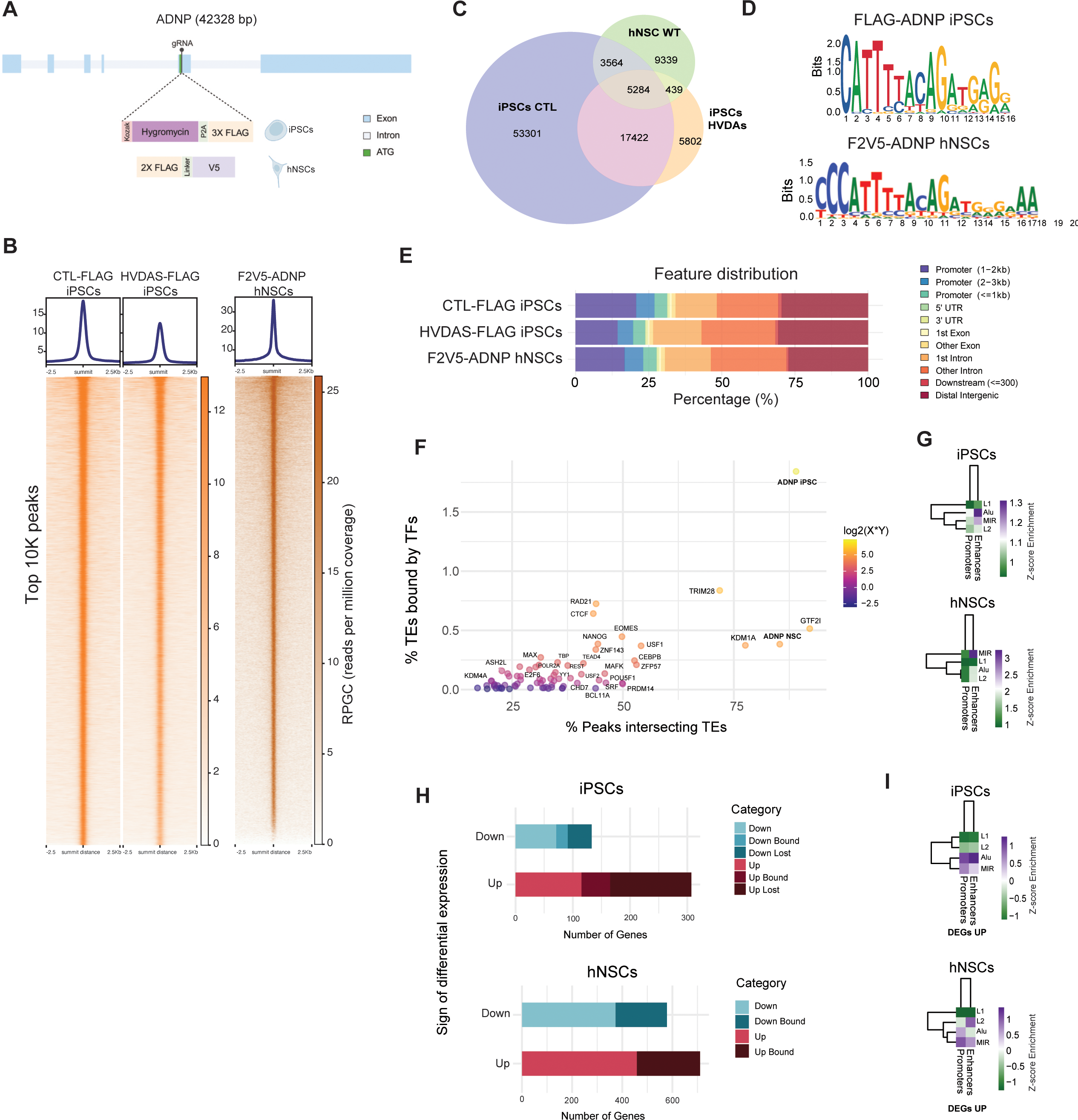
Characterization of ADNP genome binding in iPSCs and hNSCs. **A.** Schematic representation of endogenous tagging with FLAG (for iPSCs) or FLAG and V5 (for hNSCs) insertion at the N-terminus of ADNP. **B.** Heatmap of ADNP ChIP-seq enrichment in CTL-FLAG iPSCs, HVDAS-FLAG iPSCs, and F2V5-hNSCs. Each row represents a 5Kb window centered on an ADNP peak summits, sorted by ADNP ChIP signal (RPGC normalized on library size). Top 10 thousand ADNP peaks are reported. **C.** Venn Diagram showing the overlap of significant ADNP ChIP-seq peaks in HVDAS-FLAG, CTL-FLAG and F2V5-ADNP hNSCs. **D.** ADNP DNA binding motifs in iPSCs and hNSCs found *de novo* on top 10K peaks. **E.** Distribution of ADNP ChIP-seq peaks across different genomic features in CTL-FLAG, HVDAS-FLAG and F2V5-ADNP hNSCs. **F.** Scatter plot comparing percentage of TEs bound by TFs (y-axis) with percentage of TF peaks intersecting a TE (x-axis). Data for all the listed TFs refer to hESC and iPSC ENCODE-deposited ChIP-seq datasets plus our ADNP ChIP-seq performed in iPSCs and hNSCs. **G.** Heatmaps showing the enrichment of genes bound by ADNP at TEs in iPSCs (above) and hNSCs (below). The scale represents the enrichment of expressed genes bound by ADNP at TE with respect to all expressed genes bound by ADNP. **H.** Stacked barplots representing proportions of differentially expressed genes as reported in the colour legend. Bound DEGs are defined by intersecting gene regulatory regions from 4DGenome atlas of hESC and their promoters. The comparison by hypergeometric test of “DEGs” with “Bound DEGs” using expressed genes as background is significant (p=2.62e-26). The comparison of “bound upregulated genes” with “bound DEGs” is significant (p=2.15e-3, hypergeometric test). In the barplot, the “bound” groups refer to the differentially expressed genes that retain ADNP binding at regulatory regions in ADA03-FLAG, while “lost” groups refer to genes that lose ADNP binding sites in ADA03-FLAG. **I.** Heatmaps showing the enrichment of differentially expressed genes bound by ADNP at TEs in iPSCs (above) and hNSCs (below). The scale represents the enrichment of genes bound by ADNP at TE with respect to all DEGs bound by ADNP

### ADNP forms a complex with KDM1A and GTF2I to regulate gene expression

To investigate how ADNP regulates its target genes, we purified FLAG-tagged ADNP protein from nuclear extracts of F2V5-ADNP hNSCs and identified its co-purifying interaction partners by mass spectrometry (MS). Purifications from untagged wild-type hNSCs served as negative control. We identified 39 ADNP interactors of which 32 were present in all three purifications and 7 were present in two out of three purifications **(Fig.S3A, Table S3)**. We confirmed that ADNP is part of the ChAHP complex (CHD4 and HP1ɣ) also in hNSCs, and that TFIIIC interaction is observed in hNSCs as well (**Fig.3A, Fig.S3A, Table S3**)^21^. We confirmed that ADNP interacts with autism-linked POGZ and the R-loop-associated ATRX complex (ATRX, DAXX and SETDB1)^25,26^. Among the MS hits, we also observed G9a/GLP complex (EHMT1/2, WIZ, ZNF644)^27^ and 5FMC Complex (WDR18, TEX10, LAS1L, PELP1)^28^. Interestingly, we found all subunits of the LSD1 complex (KMD1A, GTF2I, ZMYM2/3/4, RCOR1/2) present in our ADNP purifications (**Fig.3A, Fig.S3A, Table S3**)^29–32^. We further focused on the LSD1 complex, as its subunits KDM1A and GTF2I show up as ADNP co-regulators of many of the same differentially expressed genes found both in iPSCs and hNSCs (**Fig.1G,H**). We confirmed the interaction of ADNP with the LSD1 complex by co-immunoprecipitation of endogenous KDM1A and GTF2I with ADNP (**Fig.3B,C**). We subsequently performed KDM1A and GTF2I ChIP-seq in wild-type hNSCs. KDM1A and GTF2I have 26,291 and 125,600 significant genome-wide binding sites respectively, and show strong enrichment in ADNP-bound regions (**Fig.3D, Fig.S3B**). We quantified the overlap between ADNP-KDM1A-GTF2I, hereafter referred to as AKG, and observed that ADNP shares the vast majority of its binding sites (14,856 or 80%) with KDM1A and/or GTF2I (**Fig.3E**), and that AKG binding sites share the same genome-wide distribution as ADNP sites (**Fig.S3E**). Because AKG significantly binds regulatory regions of DEGs (**Fig.S3F**) (∼15% of total DEGs, p=1.89e-21), we asked whether the absence of ADNP had an effect on the recruitment of KDM1A, which was the most significantly enriched TF among DEGs (**Fig.S3E**). To test this hypothesis we performed the ChIP-seq of KDM1A in the ADNP KO hNSCs (**Fig. S3D**). We found the binding motif of ADNP was less enriched among KDM1A regions in ADNP-KO hNSCs (**Fig. 3F**). Additionally, we found that a significant portion of AKG-bound DEGs lose KDM1A binding in ADNP KO hNSCs (∼29%, p=1.17e-13) (**Fig.3G**), suggesting that ADNP guides KDM1A recruitment at a subset of ADNP target genes. Focusing on genes whose promoters lose KDM1A upon ADNP KO, we observed a systematic increase in H3K4me3 (p=4,17e-3, **Fig.3H**). Given our previous observations that ADNP, KDM1A and GTF2I stand out in their percentage of target sites overlapping TEs (**Fig.2H**), we further applied K-means clustering on the whole ENCODE list of TFs to test whether ADNP binding sites are preferentially shared with KDM1A and GTF2I also in iPSCs. Indeed, we found that KDM1A and GTF2I stand out compared to all other H9 ENCODE TFs for the number of sites shared with ADNP (**Fig.S3G**). Moreover, KDM1A and GTF2I highly cluster with ADNP genome-wide positioning at promoter regions (**Fig.S3H**), suggesting that a coordinated regulatory function of AKG is also present in iPSCs.

**Figure 3.**
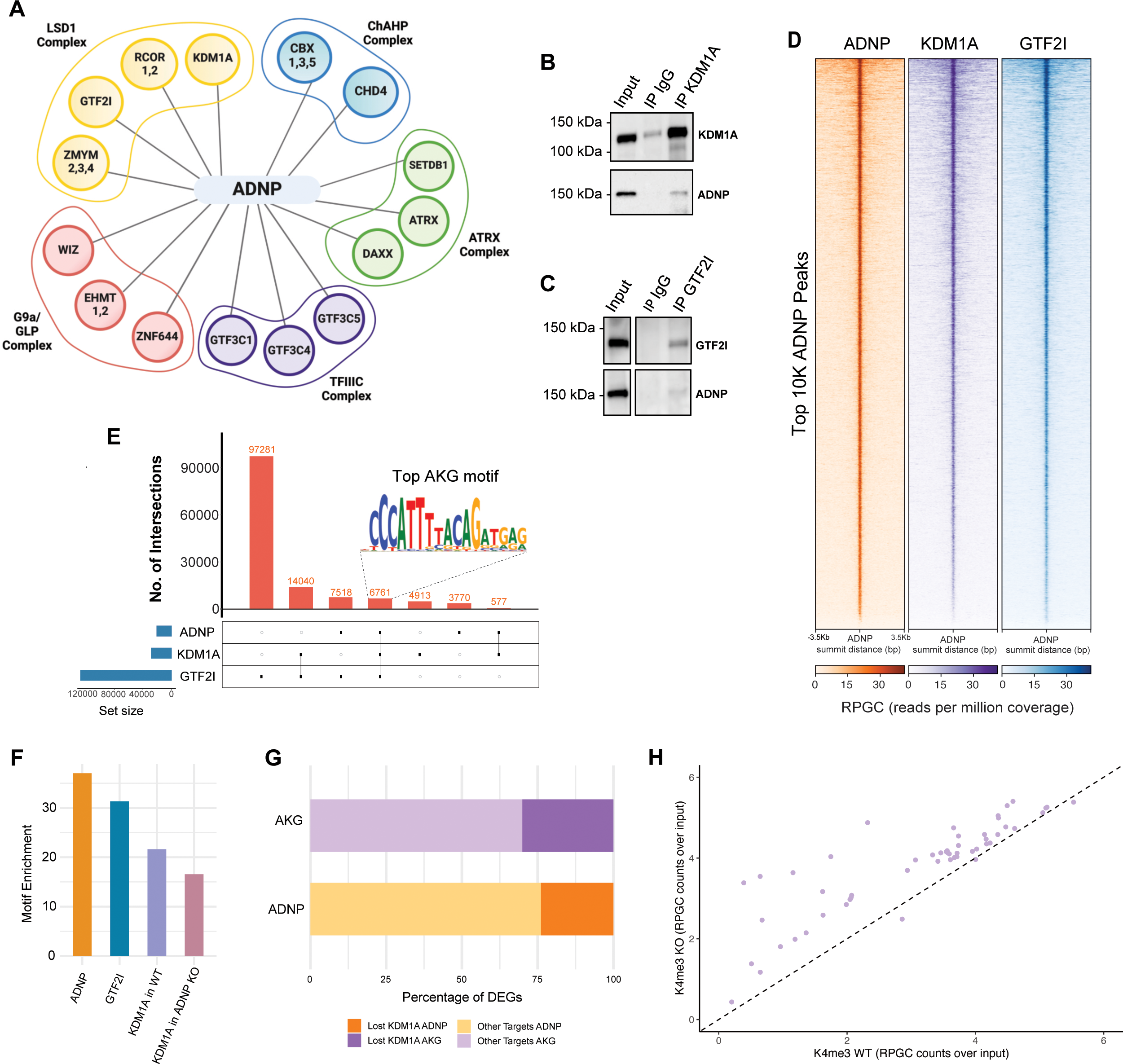
ADNP, KDM1A and GTF2I form a complex and target transposable elements in hNSCs. **A.** Schematic representation of the interactome of ADNP in hNSCs. Interactors are grouped by known complex identity and coloured accordingly. See also **Figure S3A** and **Table S3**. **B.** Western blot analysis of immunoprecipitations with KDM1A antibody or an IgG control from hNSC nuclear extract. KDM1A (upper panel) and ADNP (lower panel) are detected by KDM1A antibody and FLAG antibody, respectively. Molecular weight (MW) markers are indicated. **C.** Western blot analysis of immunoprecipitations with GTF2I antibody or an IgG control from NSC nuclear extract. GTF2I (upper panel) and ADNP (lower panel) are detected by GTF2I antibody and FLAG antibody, respectively. MW markers are indicated. **D.** Heatmap of ADNP, KDM1A and GTF2I ChIP-seq enrichments in hNSCs. Each row represents a 5kb window centered on ADNP peak summits, sorted by the TF ChIP-seq signal. ChIP-seq input signals for each TF are shown in **Figure S3B**. ChIP-seq reads have been normalized by library size (1x) as reads per million coverage (RPGC). Normalized signal legend with range is reported below each heatmap. **E.** UpSet plot indicating the number of significant peaks shared between ADNP-KDM1A-GTF2I (AKG). For the peaks shared by all three TFs, the top DNA binding motif is shown. **F.** Barplot reporting the enrichment of the ADNP motif in the top 10K peaks identified for each protein (ADNP, GTF2I, KDM1A and KDM1A upon ADNP KO respectively). Motif Enrichment was calculated with Gimmemotif v0.18 upon *de novo* motif enrichment analysis. **G.** Barplot showing the portion (%) of DEGs bound by AKG (purple) or ADNP (orange) that lose KDM1A at their promoters upon ADNP KO. **H.** Scatter plot of H3K4me3 signal in WT (x-axis) vs KO (y-axis). Both axes report hNSC ChIP-seq reads, normalized on library size (RPGC, see Methods) and scaled on the respective input data.

### ADNP mutations alter neurodevelopmental trajectories and accelerate neuronal maturation in cortical organoids

To investigate the impact of ADNP mutations on corticogenesis, we generated cortical organoids from the whole cohort of control and HVDAS iPSC lines, adopting a patterned protocol and the framework of standards that we recently contributed to establish^33^. HDVAS organoids showed an early phenotype already within the first 10 days, in terms of reduced size and irregular shape, in two independent differentiation batches (**Fig.4A**). We observed the formation of ventricles in the presence of canonical progenitor marker expression (PAX6, SOX2, Nestin) across HVDAS and control organoids in all lines (**Fig.S4A**). To probe whether the aberrant size of HVDAS organoids was linked to an abnormal proliferation of neuronal progenitors, we counted actively dividing cells by visualising the pH3 mitotic marker and found a significant reduction of dividing cells in HVDAS organoids, revealing a reduced proliferative capacity compared to controls (**Fig.4B-D**).

**Figure 4.**
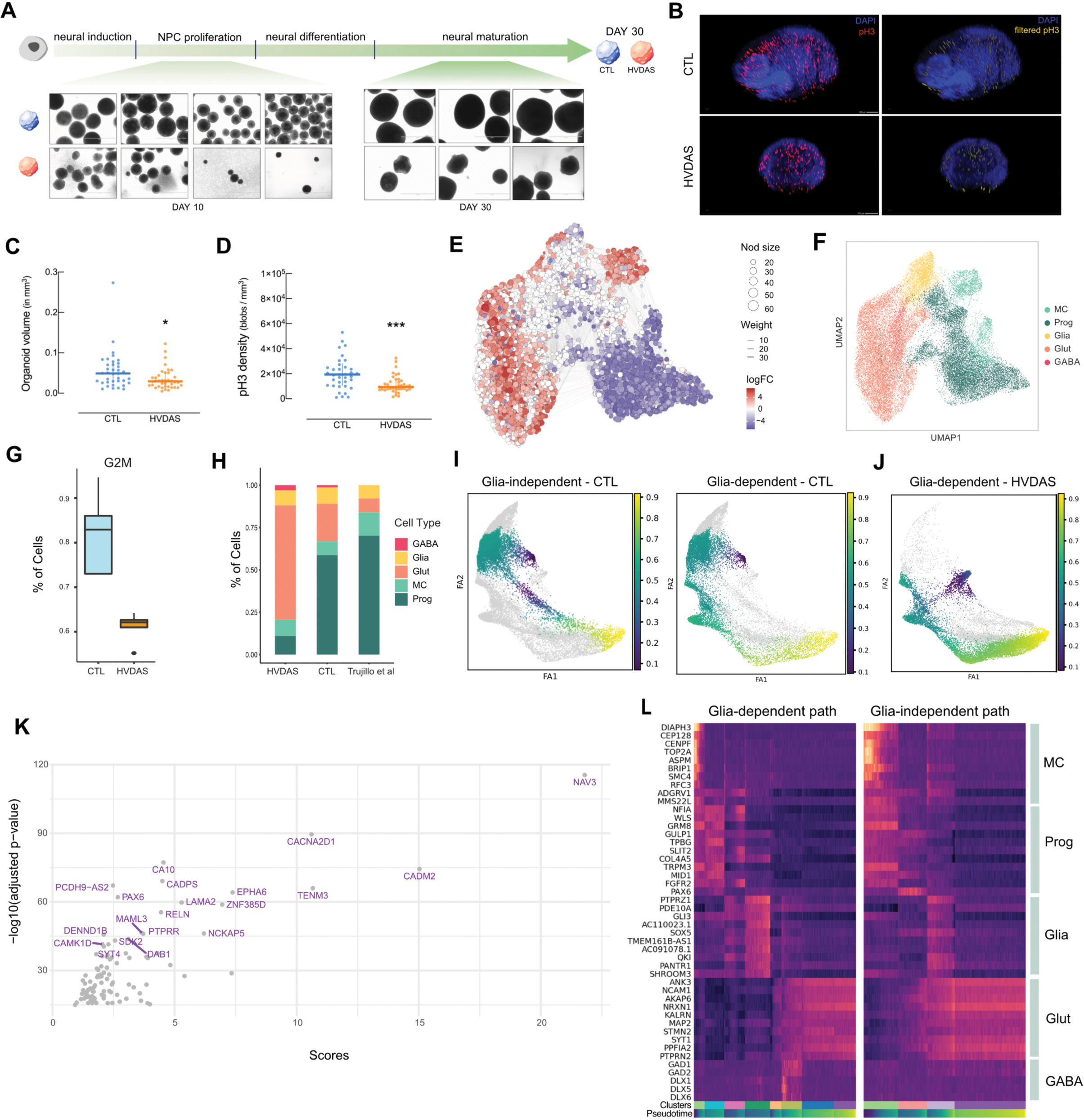
HVDAS organoids exhibit morpho-functional alterations affecting developmental trajectories. **A.** Schematic representation of cortical brain organoids generation protocol coupled with representative pictures taken along differentiation (day 10 and day 30). Pictures represent organoids derived from multiple control and HVDAS lines at indicated time points. **B.** Representative picture of cleared and immunostained control and HVDAS cortical organoids stained with DAPI and anti-pH3. The cross-section blobs identified by segmenting pH3-positive cells that are DAPI-positive are shown in yellow for both conditions. **C.** Volume measurement in control and HVDAS organoids by quantifying the signal in the DAPI channel. HVDAS n = 40, 0.02943 mm3; CTL n = 38, 0.04900 mm3; Two-tailed Mann-Whitney test * p-val = 0.0201). **D.** Phospho-H3 density measurements normalized to the size of each organoid. HVDAS n = 40, 9348 blobs/mm3; CTL n = 38, 19392 blobs/mm3; Two-tailed Mann-Whitney test *** p-val = 0.0002. **E.** Differential abundance of HVDAS cells with respect to control measured by MiloR, plotted on the UMAP of the transcriptional modality. LogFC, and Weight were calculated via miloR. Nodes are neighborhoods of cells, colored by their log fold change across conditions (CTL or HVDAS). Non-differential abundance neighborhoods (FDR >= 0.1) are colored in white, and sizes correspond to the number of cells in each neighborhood. Graph edges depict the number of cells shared between neighborhoods (Weight). The layout of nodes is determined by the position of the neighborhood index cell in the UMAP **F.** UMAP of day 30 organoids coloured by cell type (34,038 cells). MC stands for “mitotic cells”, Prog for “neural progenitors”, Glia for “glia cells” as described in Trujillo et al.^36^, Glut for “glutamatergic neurons” and GABA for “GABAergic neurons”. **G.** Boxplot reporting percentage of progenitor cells in G2M phase in CTL (light blue) and HVDAS (orange) organoids (p=8.864e-15, t-test). **H.** Barplot depicting the percentage of cells per cell type divided by genotype and compared to the reference cell atlas^36^. **I.** ForceAtlas drawgraph representing the two main paths of differentiation identified by trajectory reconstruction colored by pseudotime. On the left, CTL cells occupying the leiden clusters located on the “glia-independent” path, connecting progenitors (Prog) directly with mature neurons (farthest Glut leiden cluster). On the right CTL cells occupying the “glia-dependent” path, going from mitotic (MC) to progenitor (Prog) to glial (Glia) to early and late neurons (Glut). **J.** ForceAtlas drawgraph showing no HVDAS cells is occupying the “glia-independent” path (grey), and cells coloured by pseudotime on the “glia-dependent” path. **K.** Top 25 expressed genes in the leiden cluster 12. The score is measured via Scanpy, as a production of logFC and FDR, upon performing differential expression analysis between leiden cluster 12 and the other leiden clusters in single-cell using Wilcoxon test. **L.** Heatmaps representing the expression of marker genes for the main cell type populations identified along the trajectories, using the connections identified between leiden clusters with Paga. The two heatmaps represent expression levels scaled upon binning groups of 50 cells along the trajectory composed ordering the leiden clusters (“cluster” in the annotation reported below each heatmap). Cells in each cluster are ordered by *pseudotime* (reporting below the “cluster” annotation). The two heatmaps represent expression levels for the “glia-dependent” (left), and “glia-independent” (right) differentiation trajectories. The top markers are reported by cell type (indicated on the right side of the heatmap).

Next, to probe HVDAS neurodevelopmental alterations at molecular resolution, we performed single-cell multiomics - RNA-sequencing (scRNA-seq) coupled with single-cell assay for transposase-accessible chromatin (scATAC-seq) - on 30-days-old cortical organoids from 5 control and 5 HVDAS lines. Building on the multiplexing methodology we recently developed to increase throughput and sampling resolution, we multiplexed lines after single-cell dissociation^34^, to obtain 34,038 high quality cells. *In silico* genetic demultiplexing confirmed a balanced representation of genotypes across the entire cohort (**Fig.S4B**). We observed a clear separation between HVDAS and control cells in the uniform manifold approximation and projection (UMAP) of the RNA modality, which was confirmed by differential abundance testing with MiloR^35^ (**Fig.4E**). To understand if this separation was a result of distinct cell type abundances we applied Leiden clustering (**Fig.S4C**) to separate groups of cells based on their transcriptional similarity, and assign cell types to each cluster. Based on the expression of canonical cell type markers and retracing the expected cell types of the original organoid differentiation protocol^36^, we annotated five major populations: mitotic cells (MC), neuronal progenitors (Prog), radial glial cells (Glia), glutamatergic neurons (Glut) and GABAergic neurons (GABA) (**Fig.4F, Fig.S4D**). In agreement with the results from immunostaining, HVDAS MC clusters showed a decrease of cells in G2M (**Fig.4G**). This was consistent with the reduction of the progenitor pool, and the increase in the relative abundance of glutamatergic neurons in HVDAS organoids (p=9e-4, t-test), overall suggesting an accelerated maturation in the HVDAS samples (**Fig.4H**). To further characterise HVDAS neuronal differentiation, we ordered cells along differentiation paths and calculated diffusion pseudotime^37,38^ (**Fig. S4E**, see methods). While most cell types occupy the same space across this dimensionality reduction, mitotic cells showed a clear displacement between controls and HVDAS), with the latter showing higher expression levels of *SLC1A3*, a marker of radial glia cells (see clusters 9 and 11 in **Fig.S4F**). By inferring the progression of cells through neuronal differentiation, we found two main trajectories. We named “glia-independent” the trajectory that goes from MC to glutamatergic neurons via the progenitor pool, stemming from PAX6-expressing cells (**Fig.4I, left panel**). This path is depleted of HVDAS cells, which instead exclusively follow a second trajectory that links MCs to glutamatergic neurons, passing through early neural stem cells (Prog) and radial glia (Glia), and which we refer to “glia-dependent” path (**Fig.4I, right panel** and **Fig.4J**). The control-specific glia-independent Leiden cluster shows among the highest expressed genes markers of postmitotic neurons (NAV3), radial glia (PAX6), and early-born transient neuronal Cajal–Retzius cells and GABAergic neurons (RELN) (**Fig.4K**)^6,39–41^. The decrease in G2M vs S states in MC, and the reduced abundance of Prog populations point towards a restriction in the ability of HVDAS neural stem cells to self-renew and to a potentially faster differentiation into neurons. Moreover, the necessary passage through a more mature type of progenitors (radial glia) and the increase in glutamatergic neurons abundance suggest a general increased differentiation. Consistent with the accelerated neuronal differentiation, we noticed a larger presence of cells annotated as GABAergic neurons in HVDAS organoids compared to controls. To further characterise the GABA population, we embedded control and HVDAS transcriptomes onto a reference human cortical atlas^42^, and found that cells identified as GABAergic largely resembled medial ganglionic eminence (MGE)-derived interneurons, and that their relative abundance was doubled in HVDAS (**Fig.4SG**). Altogether, these results indicate that ADNP is an essential driver of early cortical development by controlling the timing and the trajectories of neuronal differentiation.

### Cell type-specific gene regulatory networks highlight AKG-dependent transcriptional signature in cortical organoids

We integrated chromatin accessibility with the respective transcriptome in HVDAS and CTL organoids to identify genes whose regulatory regions were preferentially accessible in each cell type. HVDAS organoids showed a higher number of markers per cell type, recalling the general tendency for gene upregulation already observed in iPSCs and hNSCs (**Fig.5A,B**), and showed a higher distinction between cell types. Nevertheless, we could not find crucial neuronal fate determinants such as PAX6, which was present only in the progenitors of control organoids. Next, we performed differential expression analysis on each cell type, by comparing controls and HVDAS organoids in pseudo-bulk^43^. We analysed differential TF activity between CTL and HVDAS in each cell type, and found increased accessibility of genomic regions bearing OCT and SOX motifs in progenitors, glia and GABAergic neurons (**Table S4**) (**Fig.S5A**). Mitotic cells are the only population that form two separate clusters either populated by CTL or HVDAS cells (**Fig.4E,F**). Accordingly, we focused on the regulatory regions of mitotic cells DEGs harbouring an AKG motif (p < 0.001) to evaluate a direct contribution of the complex in this transcriptional outcome. GO enrichment on DEGs putatively regulated by AKG, found *neurogenesis* as the most represented category (**Fig.5SB**). In addition, the majority of neurogenesis DEGs harbouring the AKG motif were upregulated in mitotic cells of HVDAS organoids (**Fig.S5C**). The same genes were steadily dysregulated across cell types (**Fig.S5D**). This points to sustained activation of the neurogenesis program in HVDAS via AKG-regulated genes. Among the genes putatively under AKG control and systematically upregulated across HVDAS cell types, we identified neurogenesis-related *AUTS2*, a gene implicated in numerous neurological disorders, including ASD. Its increase in expression was coupled with a higher activity of an HVDAS-specific co-accessible distal CRE containing the AKG motif (**Fig.5C**). Cell type-specific DEGs were significantly enriched in AKG targets, in particular DEGs found in the three cell types constituting the *glia-dependent* path (MC-Glia-Glut) (**Fig.S5E**). We applied CellOracle^44^ to generate cell type-specific gene regulatory networks (GRNs) and identify differentially active TFs that significantly explain the differential expression observed in each cell type, independently of AKG (**Fig.5D**). To estimate the contribution of each of these master regulators to the previously observed cell type composition defects we performed *in silico* KO in HVDAS cells. This allowed us to identify those TFs capable of increasing the progenitor pool while reducing glutamatergic neurons abundance (**Fig.5E**). Notably, several perturbations, including all the AKG targets, rescued the overall cell type composition by also reducing GABAergic populations (**Fig.S5F**).

**Figure 5.**
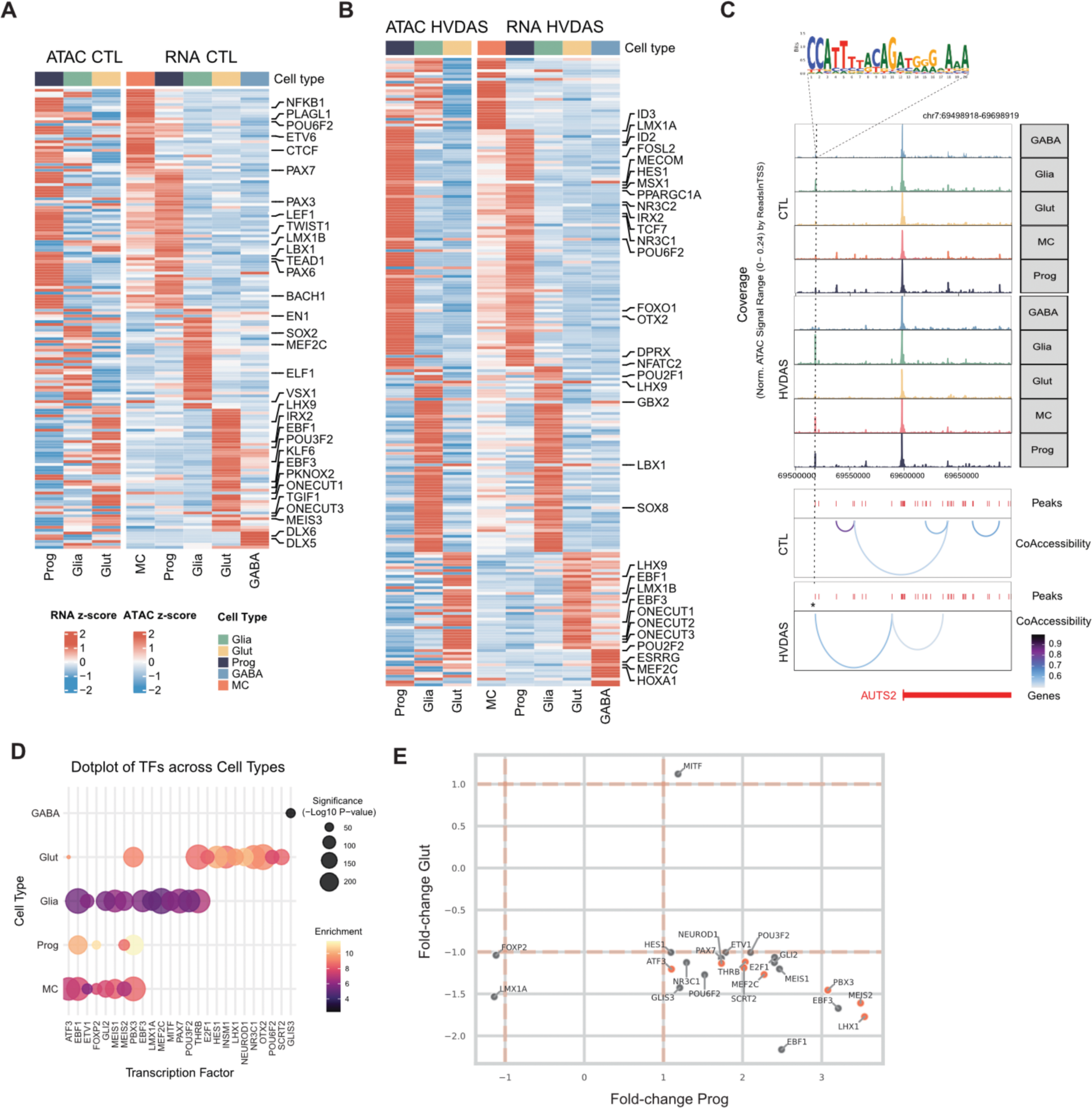
AKG-dependent transcriptional dysregulation in cortical organoids. **A.** cell type-specific markers identified by integration of ATAC and transcriptomic modalities via multiomics profiling of control cortical brain organoids. Heatmaps represent z-scores of both ATAC and RNA signals across cell types. All chromatin regions and genes associated to a transcription factor are labeled on the right side of each heatmap. cell types are indicated on the bottom (x-axis). **B.** cell type-specific markers identified by integration of ATAC and transcriptomic modalities via multiomics profiling of HVDAS cortical brain organoids. Heatmaps represent z-scores of both ATAC and RNA signals across cell types. For each couple of chromatin accessibility and gene expression signals, transcription factors are labeled on the right side, and cell types are indicated on the x-axis. **C.** Exemplary tracks of an AKG target involved in neurogenesis (*AUTS2*) whose co-accessibility pattern changes in HVDAS. Pseudobulk ATAC coverage is reported for cell type, first in controls (above), and then in HVDAS (below). Chromosomal location is reported on top-right. The location of the AKG motif is indicated by a dashed line and a motif logo. Co-accessibility patterns are depicted for controls and HVDAS separately, to highlight the gain of accessibility at AKG binding site. The windows are centered on the *AUTS2* transcription start site. Peaks found across samples and replicates (at least two multiomic sequencing runs) are indicated as red bars. Co-accessibility signal is reported as blue links. **D.** Dotplot of enrichment and significance of regulatory activity by TFs identified in gene-regulatory networks (GRN) of each cell type. Each TF reported on the x-axis is differentially expressed. Dot color refers to the enrichment of targets of each TF in the GRN with respect to the differential expression analysis of each cell type. Dot size refers to significance (-log10 p-value) calculated with hypergeometric test comparing targets of each TFs in the GRN of a certain cell type with the DEGs of the same cell type, using expressed genes in said cell type as background. **E.** Scatter plot of the effect of virtual KO measured for each master regulator in terms of relative cell type abundance before and after KO in HVDAS lines via CellOracle. Fold-change of progenitors (Prog) abundance after KO is reported on the x-axis. Fold-change of glutamatergic neurons (Glut) abundance upon KO is reported onthe y-axis. Red dots refer to TFs whose KO increase Prog abundance, while decreasing Glut and GABAergic neurons abundance.

Then, we extracted the MC-specific network of *neurogenesis* genes regulated by AKG (**Fig.S5G**) and found three upregulated TFs (GLI2, ETV1 and PBX3), which were able to rescue the accelerated maturation phenotype in the *in silico* screening, suggesting that their derepression may be a direct cause of *neurogenesis* dysregulation in HVDAS organoids. Next, we focused on glutamatergic neurons, enrichment for DEGs bearing an AKG motif at their regulatory regions was *behaviour*, and the majority of DEGs in this category were upregulated (78%), including *PBX3*. Of note, among upregulated genes we found *GAD1*, a marker of GABAergic neurons, which is both bound by ADNP in control iPSC and upregulated in HVDAS iPSCs. Altogether, these results show that the altered expression of specific target genes under the control of AKG contributes to explaining the disrupted GRNs and cell-fate decisions observed in HVDAS organoids.

## DISCUSSION

The complex genetic architecture of neurodevelopmental disorders poses two complementary challenges for mechanistic dissection and mental health care alike. The first is to locate at which levels convergences lie that can best explain how different NDD-associated mutations or polygenic risk loadings bring about shared phenotypes. The second is to probe those levels to dissect how NDD-causative genes also display specific phenotypic spectra and which gene-specific early developmental milestones can thus inform personalized prognosis. As shown in the recent exhaustive phenotypic comparison across ADNP-, CHD8- and DYRK1A-NDD^45^, beyond the shared association with ASD and ID it was indeed possible to outline gene-specific phenotypic vulnerabilities, with oppositional features overrepresented in HDVAS. At the current level of clinical granularity, though, early developmental milestones showed limited predictive value, underscoring the importance of probing neurodevelopmental antecedents both at earlier timepoints, as now enabled by organoid technology, and at high molecular resolution. Focusing on HVDAS as an NDD paradigm, our work provides the following insights of broad relevance for the modelling of such conditions.

First, we found ADNP to be particularly enriched at enhancers and promoters falling within primate-specific Alu elements, which represent 27% of the primate-specific TE-derived TFs binding sites^46^, suggesting a concomitant evolutionary rewiring of ADNP activity. As expected for loss-of-function pathogenic mutations of a chromatin factor that we found to be predominantly repressive, genes whose expression was affected by genome-wide loss of ADNP at regulatory regions were mostly upregulated. A significant portion of those regions was found within TEs, meaning that ADNP modulates gene expression by repressing TE-situated regulatory regions^47^. Alu elements have been long recognized for their key role in shaping the primate brain, with their retrotransposition events estimated to have occurred at 15-fold higher rate in the human, chimpanzee, and bonobo lineage (compared to other great apes) and at double rate in humans vis a vis chimpanzee and bonobo^48–50^. Moreover, there is a growing list of neurodevelopmental and neurodegenerative conditions with an established or candidate causative role for deleterious Alu activity^48^. Our data on ADNP show how a single dosage-sensitive factor brings about Alu-mediated pathogenicity affecting brain functional domains, providing another example of the intricate relation between NDDs and modelling of human evolution^51–53^.

Second, we retrieved a shared ADNP DNA-binding motif from both iPSCs and hNSCs, which, in contrast with what was previously reported in mouse embryonic stem cells, does not match that of CTCF^20^. Comparing ChIP-seq from ADNP and CTCF we found very limited occupancy overlap, pointing to a species-specific switch in TF genome-wide binding sites between human and mouse^46,54,55^. Together with the major impact on Alu-containing regulatory regions, these findings invite caution in extrapolating conclusions from more distantly related species, including mice, especially with the aim of understanding quintessentially human conditions.

Third, consistent with the role of ADNP in transcriptional repression, our protein interactome in hNSCs, revealed that ADNP mostly binds co-repressors, such as the LSD1 complex^56^, the G9A/GLP complex^57^, as well as ChAHP and TFIIIC complexes, that were previously described^19,21^. We focused on the interplay with KDM1A and GTF2I, which we refer to as AKG complex, because i) *ADNP*, *KDM1A* and *GTF2I* featured together in both iPSCs and hNSCs master gene regulator analysis, ii) KDM1A and GTF2I cluster together with ADNP in genome-wide promoter occupancy of pluripotent stem cells, and iii) they all have an exceptionally high fraction of their binding sites in TEs, overall pointing to converging mechanisms of gene regulation. As the LSD1 complex is recruited by sequence-specific transcription factors to remove the activating H3K4me2 and H3K4me1 marks from gene-regulatory regions^58,59^, KDM1A plays a fundamental role in enhancer silencing in multiple cell lineages^60^. We biochemically validate the interaction of ADNP, KDM1A and GTF2I, and find a strong overlap in genome-wide binding patterns and motifs. hNSCs depleted of ADNP displayed a loss of KDM1A binding at the promoters of a significant subset of upregulated genes, suggesting that ADNP is required for the recruitment of KDM1A to such targets. Indeed, ADNP-bound promoters that lose KDM1A significantly enriched for regions gaining H3K4me3 in ADNP KO hNSCs compared to wild-type. Thus, our data indicate that ADNP recruits KDM1A to specific targets to repress gene expression at regulatory TEs. Of note, we had previously shown that KDM1A recruitment by GTF2I in 7dupASD (the ASD syndrome caused by microduplication of 7q11.23) underlies GTF2I dosage-dependent gene repression^29^, warranting the use of KDM1A inhibitors to rescue core ASD proxy symptoms in murine models harboring GTF2I duplications^61^. The identification of an ADNP-GTF2I-KDM1A complex, uncovers thus a potential mechanistic convergence between these two paradigmatic ASD conditions.

Fourth, our single-cell multiomic profiling of HVDAS organoids positions ADNP mutations in the context of the imbalances in temporal trajectories that are being recognized as a fundamental level of dysregulation in NDDs^62–66^. Specifically, we showed a large disproportion in the abundance of neural progenitors and glutamatergic neurons, with HVDAS showing a higher percentage of the latter. This accelerated neuronal maturation phenotype, with the presence of GABAergic neurons nearly exclusive to HVDAS organoids, is in line with our previous findings in CHD8+/-organoids^67^, indicating a trajectory-level cellular antecedent across disorders. Developmental trajectory analyses showed that control organoids follow two main differentiation patterns: one that passes through the radial glia, and an earlier one that passes through a *SLC1A3-*/*GLAST-* progenitors population. The latter trajectory appears to be completely depleted in HVDAS organoids, with the remaining one generating neurons mainly expressing migratory and upper-layer neuronal markers. This is in line with a perturbation screening showing that ADNP deficiency leads to the disproportional formation of upper-layer neurons in telencephalic organoids^68^. In addition, a conditional knock-out of ADNP in mice showed that ADNP is critical for driving the expansion of upper-layer cortical neurons^69^. Put into context, ADNP seems thus to relate to a set of chromatin factors such as EZH2, EHMT1, EHMT2 or DOT1L which establish cell-intrinsic timing during human neuronal differentiation, and whose perturbations cause acceleration of neuronal maturation^70,71^. In this light, we investigated the disrupted GRNs underlying premature neurogenesis in HVDAS organoids and found that neurogenesis genes that are normally bound by AKG are de-repressed in HVDAS mitotic cells. This suggests that the failure of AKG to timely repress such genes causes premature activation of the pro-neural fate program and subsequent aberrant neurogenesis. Finally, we identified the AKG targets determining differentiation biases in HVDAS organoids. *In silico* knock-out of such master regulators univocally corrected the aberrant progenitor, glutamatergic and GABAergic imbalances, at once. Notably, *PBX3* appeared to be a key driver of neurogenesis, normally repressed by AKG in mitotic cells and part of the constitutively upregulated *neurogenesis* transcriptional program, positioning it as one of the earliest AKG targets driving HVDAS differentiation defects. *GAD1* is another relevant example, as its regulation influences the production of GABAergic neurons, and has been associated with schizophrenia and ASD^63,73^. We showed that *GAD1* is a target of ADNP, repressed both in iPSCs and cortical organoids suggesting that AKG also contribute to GABAergic interneuron differentiation during the early stages of corticogenesis.

Together, this first patient-specific modelling of HDVAS uncovers ADNP as a transcriptional repressor exhibiting broad functional convergence with the core of chromatin gatekeepers controlling neurodevelopmental tempo, paving the way to a trajectory-level inference of NDDs convergences and specificities.

---

**Figure S1.**
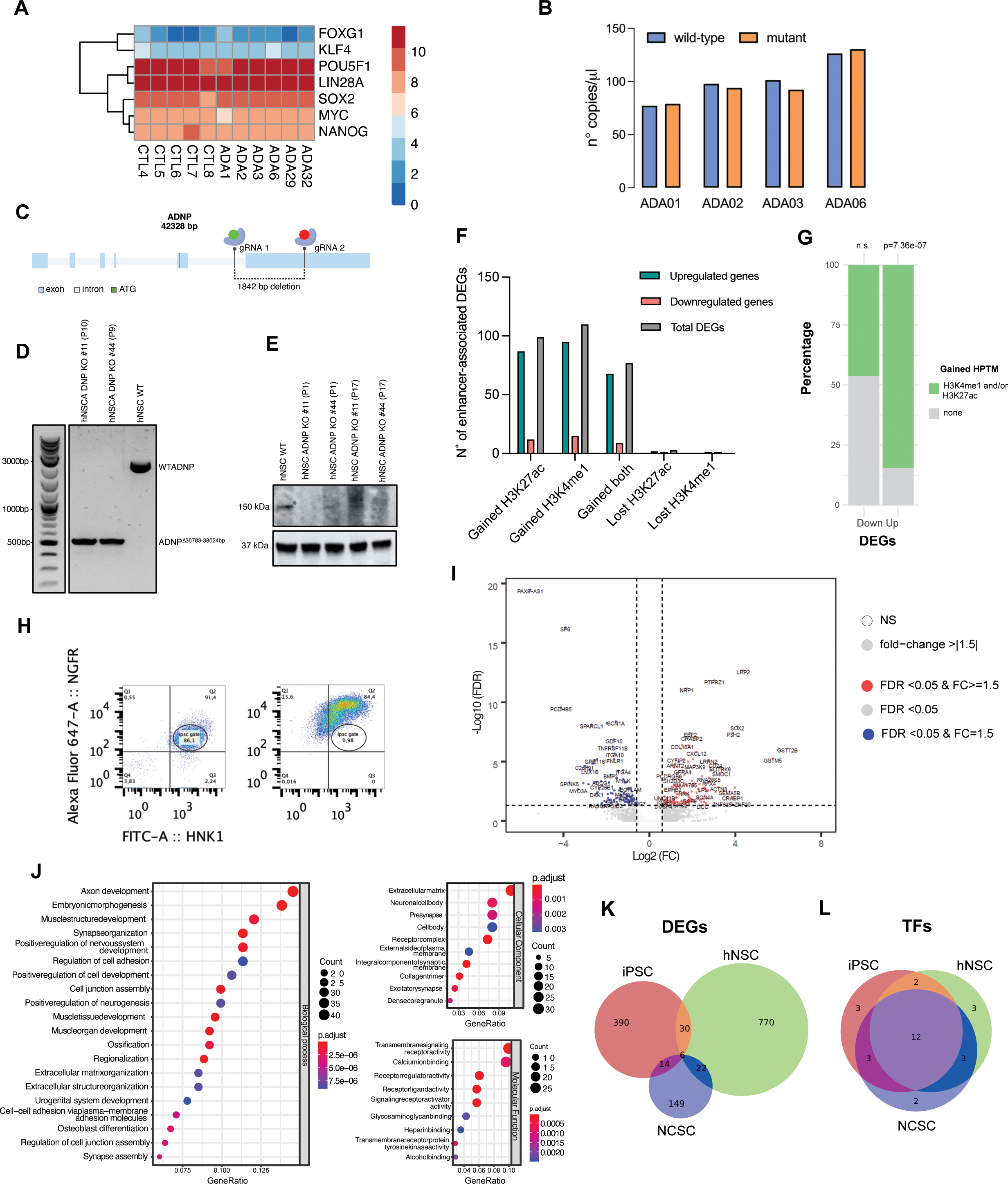
Gene dysregulation in HVDAS iPSCs and ADNP-KO hNSCs. **A.** Heatmap showing logTMM expression of pluripotency markers in the whole cohort of iPSC samples **B.** dPCR quantification of allele-specific amplicons for the indicated patient-derived iPSC samples. wild-type and mutant alleles are respectively colored in blue and orange. **C.** Schematic representation of ADNP knock-out editing strategy. ADNP gene structure is reported. Cutting target sites for the two gRNAs are indicated upstream (green) exon 5 and after 1842 bp (red) (see **Table S5**). **D.** PCR amplification of the ADNP KO regions for two different correctly edited clones versus the WT sample (see **Table S5**). **E.** Western Blot showing the absence of ADNP protein in the edited clones versus the WT sample. **F.** Barplot showing the intersection of DEGs (sub-sampled in upregulated and downregulated) with annotated enhancers (4DGenome repository), and the gain or loss of H3K27ac, H3K4me1 or both. **G.** Percentage of downregulated (Down) and upregulated (Up) genes that show a gain in H3K27ac and/or H3K4me1 at their annotated enhancers (green). Significance of genes that gained either or both marks contributing to the two categories is reported above each bar. **H.** Gating strategy for the identification of NCSCs, based on two markers: NGFR and HNK1 **I.** Volcano plot reporting the differentially expressed genes in HVDAS iPSC with respect to controls. The X-axis represents log2(fold change) and the Y-axis represents -log10(FDR). **J.** GO terms for NCSC RNA-seq. ‘Gene ratio’ indicates the portion of total DEGs in the given GO term. Dot size (Count) represents the number of DEGs in each category. Dot color represents significance (p-adjusted). GO aspects (BP, CC and MF) are indicated on the right of each plot in a grey box. **K.** Proportional Venn diagram showing the number of DEGs shared across HVDAS iPSCs, ADNP KO hNSCs, and HVDAS NCSCs. **L.** Proportional Venn diagram showing the number of TFs shared across HVDAS iPSCs, ADNP KO hNSCs, and HVDAS NCSCs based on each independent master regulatory analysis.

**Figure S2.**
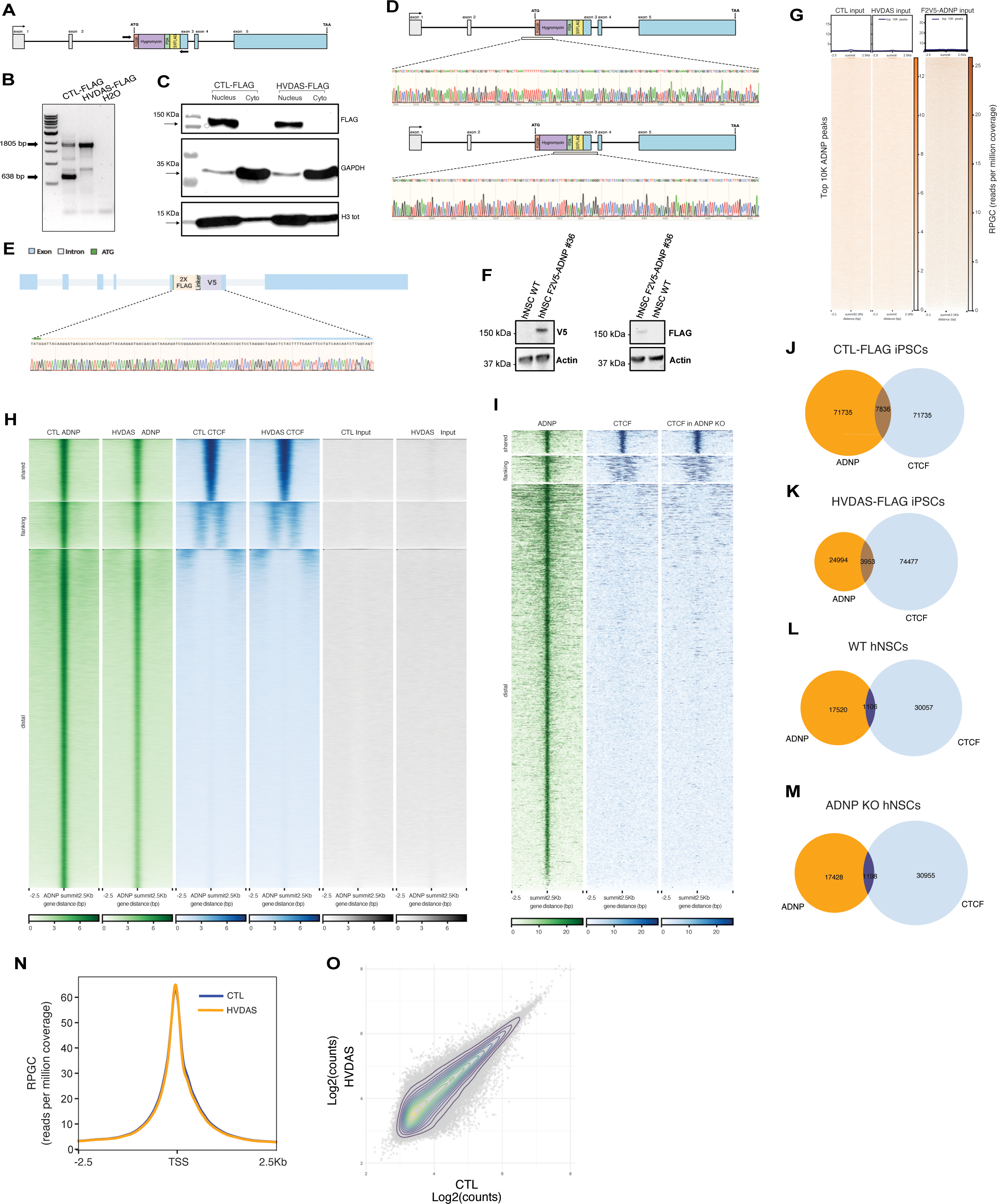
Characterization of ADNP genome binding in iPSCs and hNSCs. **A.** PCR amplification strategy at the ADNP locus for the screening of homozygous versus heterozygous FLAG-iPSC edited clones (see **Table S5**). **B.** PCR products for the selected heterozygous clone (arrows in CTL-FLAG iPSCs) and homozygous clone (arrow in HVDAS-FLAG iPSCs) resulting from edited iPSC HVDAS line. Sanger-seq results show the correct in-frame CRISPR editing of the FLAG insertion in iPSCs. **C.** Western Blot showing the differential abundance of FLAG-ADNP in cellular compartments upon nuclear/cytoplasm fractionation of the edited iPSCs line (see Methods). **D.** Sanger-seq results show the correct in-frame CRISPR editing of the FLAG insertion in iPSCs (see **Table S5**). **E.** Sanger-seq results show the correct in-frame CRISPR editing of the FLAG insertion in hNSCs (see **Table S5**). **F.** Western Blot showing the detection of hNSC tagged-ADNP line using either V5 antibody (left panel) or FLAG antibody (right panel). **G.** Heatmap of ADNP ChIP-seq input in CTL-FLAG iPSCs, HVDAS-FLAG iPSCs and F2V5-hNSCs centered on ADNP peak summits. **H.** Heatmap of ADNP (green) and CTCF (blue) ChIP-seq enrichment in CTL-FLAG iPSCs and HVDAS-FLAG iPSCs samples, with related input signals. Each row represents a 5Kb window centered on ADNP peak summits, sorted by the CTCF ChIP signal. Peaks are grouped by distance with respect to ADNP sites: shared (overlapping peaks), flanking (+/-1000 bp) and distal (> 1000 bp). **I.** Heatmap of ADNP (green) and CTCF (blue) signals at ADNP summits at shared, flanking and distal sites in hNSCs. **J.** Venn diagram showing the overlap of ADNP-FLAG and CTCF ChIP-seq peaks found in CTL-FLAG iPSCs. **K.** Venn diagram showing the overlap of ADNP-FLAG and CTCF ChIP-seq peaks found in HVDAS-FLAG iPSCs. **L.** Venn showing the overlap of ADNP and CTCF ChIP-seq peaks found in WT hNSCs **M.** Venn showing the overlap of ADNP and CTCF ChIP-seq peaks found in ADNP KO hNSCs **N.** Aggregate normalized ATAC-seq signal (RPGC) for the whole cohort of control and HVDAS iPSCs, centered on TSS. **O.** ATAC-seq correlation analysis performed on the whole cohort of control and HVDAS iPSCs.

**Figure S3.**
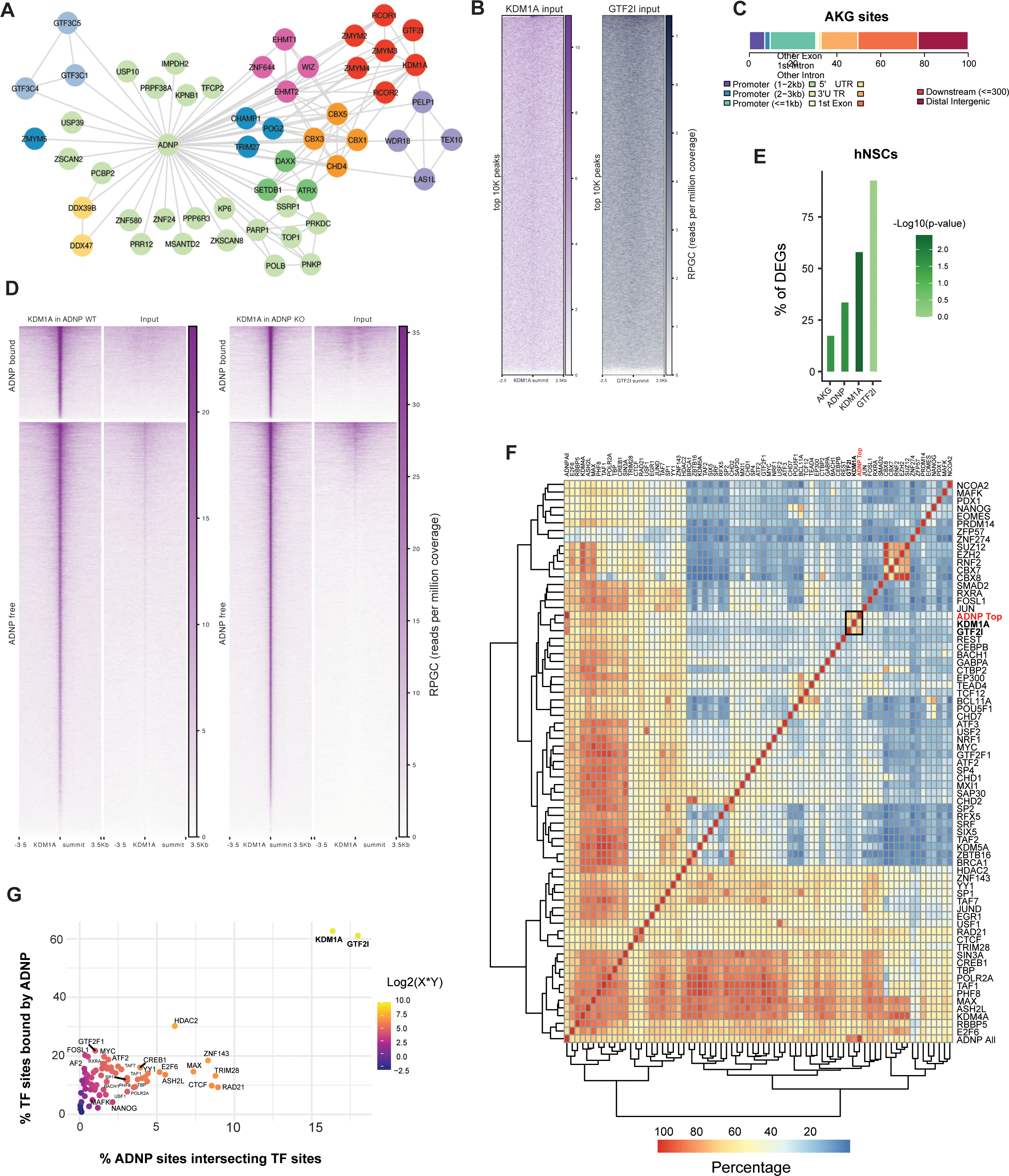
ADNP, KDM1A and GTF2I form a complex and target transposable elements in hNSCs. **A.** Complete interactome of ADNP in hNSCs derived from ChIP-MS. Nodes are colored based on their function or complex association. red: GTF2I/KDM1A complex, orange: ChAP complex, pink: EHMT1/2 complex, blue: POGZ interactors, light blue: TFIIIC complex, dark green: ATRX complex. Known physical interactions excluding ADNP were derived from Stringdb and reported. **B.** ChIP-seq input signal for KDM1A and GTF2I in WT hNSCs. Each row represents a 5Kb window centered on KDM1A and GTF2I peak summits, respectively. **C.** Distribution of peaks shared by ADNP-KDM1A-GTF2I (AKG) across different genomic features with respect to protein-coding genes. **D.** Heatmap of KDM1A ChIP-seq signal in ADNP WT and ADNP KO hNSCs. Each row represents a 5Kb window centered on peak summits, sorted by the KDM1A ChIP signal. Peaks are divided into ADNP-bound (top), and ADNP-free (bottom) based on effective ADNP binding in the WT line. **E.** Percentage of DEGs in hNSCs bound by individual AKG components or the whole complex. Significance value for each TF binding is reported. **F.** Heatmap representing the percentage of peaks found between pairs of ENCODE TFs, located within +2500bp and −2500bp from the TSS of protein-coding genes. A red square highlights a cluster made by the top 10,000 peaks of ADNP (“ADNP Top”) and peaks of KDM1A and GTF2I (in bold). All ADNP peaks are indicated as “ADNP All” in the bottom row of the heatmap. **G.** Scatterplot representing the percentage of ADNP peaks contained in ENCODE TF binding sites of H9 hESC (y-axis) and percentage of ENCODE peaks contained in CTL-FLAG iPSCs ADNP binding sites. Dots are colored by log2(%X*%Y).

**Figure S4.**
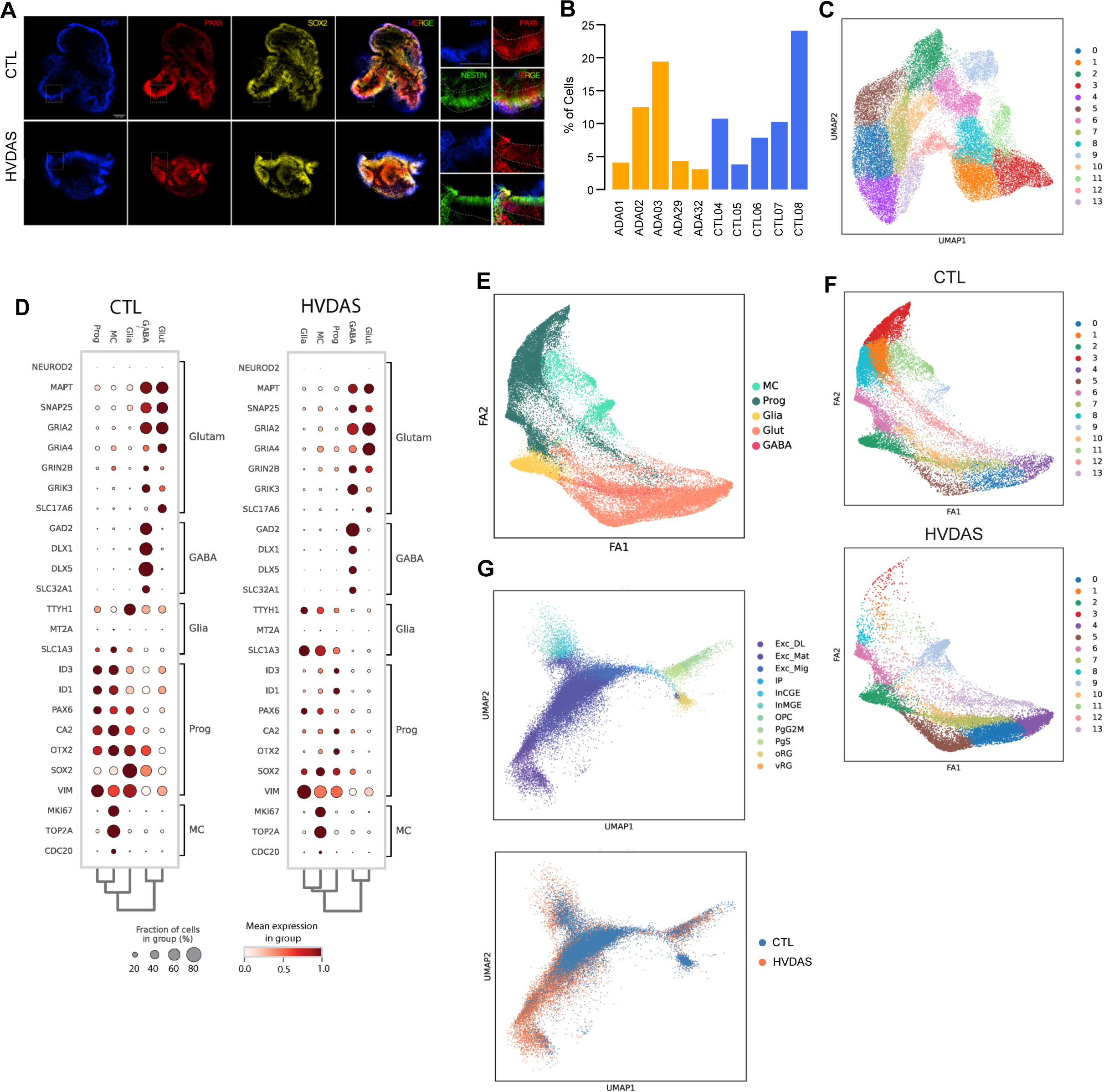
HVDAS organoids exhibit morpho-functional alterations affecting developmental trajectories. **A.** Representative picture of cleared and immunostained control and HVDAS cortical organoids stained with DAPI, anti-PAX6, anti-SOX2 and anti-Nestin. The portion of the organoids enclosed in the dashed square is magnified in the quadrants located in the right-most part of the figure. **B.** Bar plot representing the relative abundance of each individual on total number of cells in our multiome dataset. HVDAS lines are indicated in orange and control lines in blue. y-axis is expressed as percentages summing to 1. **C.** Leiden clustering reported on UMAP dimensionality reduction of the transcriptional modality of our multiome dataset, using resolution = 0.5. **D.** Dot plot showing the fraction of cells expressing marker genes for each major annotated neuronal cluster. Fraction of cells is represented by dot size, and mean expression values by color. CTL and HVDAS cells are respectively reported on the left and on the right to appreciate genotype-specific expression levels. **E.** Drawgraph representation of major cell types found in the transcriptome. Cycling progenitors (i.e. mitotic cells, MC), early neural progenitors (Prog), radial glia (Glia) and glutamatergic neurons (Glut) derive from joining leiden clusters. GABAergic neurons (GABA) were identified using a scoring function (scanpy.tl.score_genes) using GAD1/2,DLX1/5,SLC32A1 and DLX6-AS1 markers (see Methods) within the Glut population. **F.** Drawgraph representation of Leiden clusters for CTL (above) and HVDAS (below) cells to appreciate the depletion of cells per leiden cluster. **G.** UMAP representation of the transcriptional modality of controls and HVDAS organoids cells transferred by ingestion on fetal prefrontal cortex (PFC) data from Badhuri et al^42^. Upper UMAP is colored by cell type attributed to PCF data, and transferred to organoid data. Cells in the lower UMAP are colored by genotype, showing HVDAS populating inhibitory and more mature neuronal populations. Cell type labels refer to annotations derived from^74^, as described in the methods. vRG and oRG refer to ventral- and outer-Radial Glia; PgS and PG2M refer to S-phase and G2M-phase mitotic progenitors, respectively; OPC refer to Oligodendrocyte Progenitors Cells; InMGE and InCGE refer to interneurons of Medial- and Caudal-Ganglionic Eminence, respectively; IP refer to Intermediate Progenitors; Exc-Mig, Mat and DL, respectively refer to Exitatory Migrating-Maturing- and Deep Layer-neurons.

**Figure S5.**
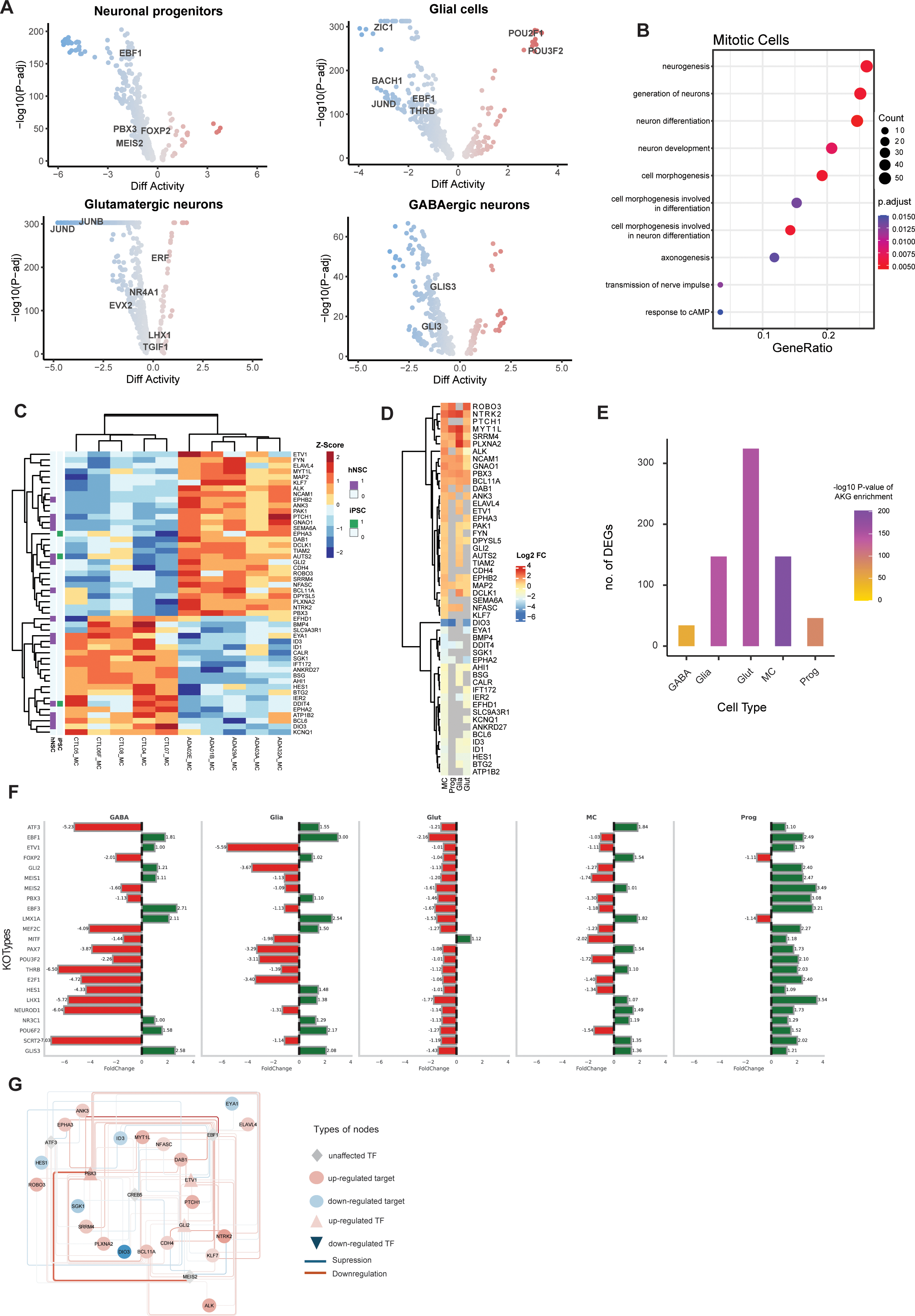
AKG-dependent transcriptional dysregulation in cortical organoids. **A.** Volcano plots of TFs differential activity measured by comparing HVDAS with control lines in the chromatin accessibility modality in each cell type using ChromVar. Labels are reported for TFs differentially expressed in HVDAS with respect to controls in the same cell type. **B.** Dotplot of Biological Process component of GO enrichments performed on differentially expressed genes regulated by AKG in mitotic cells. Dot size represents number of genes differentially expressed in the enriched category. Dot color represents significance (-log10(p-adjusted)). **C.** Heatmap of *neurogenesis* gene expression in mitotic cells measured in pseudobulk (z-scores). Each row (y-axis) is a gene. Gene names are annotated on the right, binding of the gene by ADNP in iPSC (green) and hNSC (purple) is reported on the left. Each column reports the values for each individual. Both axis are clustered through hierarchical clustering. **D.** Heatmap of log2FC for *neurogenesis* genes (y-axis) found differentially expressed in mitotic cells, measured in each cell type (x-axis). **E.** Barplot representing the number of genes found differentially expressed in each cell type, by comparing HVDAS with control cells in pseudobulk. Bars are colored by significance (-log10 p-value) based on the amount of AKG targets measured with hypergeometric test. **F.** Barplot of fold-changes calculated upon *virtual* KO of master regulators identified for each cell type. The effect of each TF KO (y-axis) is measured as a fold-change in terms of relative increase in cells for each cell type (x-axis) in HVDAS before and after the KO (% in each KO / % in unperturbed HVDAS). **G.** Gene-regulatory network belonging to *neurogenesis* category, that is affected in HVDAS cycling progenitors. The GRN has been generated by the integration of transcriptomic and ATAC modalities via CellOracle. All colored nodes are direct targets of AKG. Grey square nodes represent TFs that are not under direct control of AKG but directly bind dysregulated target regulatory regions (see figure legend).

## ACKNOWLEDGEMENTS

This work was supported by grants from the European Research Area Network of European Funding ERA-NET (ADNPinMED), and E-Rare (IMPACT). M.F.P. has benefited from the H2020 Marie Skłodowska-Curie Actions, Grant/Award Number: 765966. M.A. de P. was supported by an ALW-open program grant (ALWOP.372) from NWO. A.V. was supported by Telethon grant GP19226 granted to G.T and RE-MEND (101057604). This work was supported, in part, by US National Institutes of Health (NIH) grant R01MH101221 to E.E.E. E.E.E. is an investigator of the Howard Hughes Medical Institute. We thank the HVDAS patient community for the interest in our work and in particular the families that donated fibroblasts for research purposes. Susanne E. Boonen and Terri McVeigh are acknowledged for helping to collect patient materials.

## AUTHOR CONTRIBUTION

L.R. designed and performed all the experiments in iPSCs, NCSCs and organoids, and prepared the manuscript. A.V. analysed all the experiments and prepared the manuscript. M.P.A. designed, performed and analysed all experiments in ADNP KO hNSCs, performed F2V5-ADNP, KDM1A and GTF2I ChIPs and prepared the manuscript. L.M. generated the ADNP-F2V5 hNSC cell line and performed ADNP purifications. M.D. provided support in experiments with hNSCs and performed immunoprecipitations. D.H.W.D. and J.D. performed mass spectrometry analyses. F.D. conducted the analyses on ADNP ChIP-seq datasets, and contributed to the multi-omics analyses. V.F. contributed to the implementation of multi-omics data analysis. M.F.P. contributed with the standardisation of the organoid differentiation protocols and performed the stainings. M.G. and D.P. set up the cohort of iPSCs through reprogramming of patient-derived fibroblasts. S.T. contributed to the design of the iPSCs CRISPR engineering. M.M. helped in the interpretation of sequencing data from patient-derived cell lines. E.T. performed the library preparation for RNA-seq experiment. E.E.E. provided patient-derived fibroblasts upon clinical evaluation. C.E.P. provided collaborative support in the interpretation of data coming from patents’ cell lines. F.K. and A.V.D. provided patient-derived fibroblasts upon clinical evaluation and contributed to the design of the study. G.T. supervised the experimental design and analyses, contributed interpreting results and provided funding. G.T. L.R. and A.V. conceived the experiments on patient-derived lines. R.A.P. supervised the experimental design and analysis, contributed interpreting results and provided funding.

## DATA AVAILABILITY

Supplementary data can be found at Zenodo link : 10.5281/zenodo.14944874, and further information and code will be available upon request.

## MATERIAL AND METHODS

### Cell reprogramming

Skin fibroblasts from HVDAS patients were received from Prof. Frank Kooy (Antwerpen Universiteit, Belgium) and Prof. Even Eichler (University of Washington). Reprogramming was performed using non-integrating, self-replicating mRNAs as previously described^75^ (Stemgent, 00-0071). All control lines were reprogrammed starting from healthy donor biopsies; CTL08A was the only line purchased from the Wellcome Trust Sanger Institute and was the only reprogrammed using the Sendai virus (CytoTune-iPS 2.0 Sendai Reprogramming Kit; Thermo Fisher Scientific, A16517). The genetic and clinical details regarding mutant hiPSCs used in this work are summarised in **Table S1**.

### Cell Culture

hiPSC were cultured in TeSR-E8 medium (Stemcell technologies, 05990) supplemented with penicillin-streptomycin (P/S, 100 U/mL; Thermo Fisher Scientific, 15140-122), with daily media change, at 37 °C, 5 % CO2 and 3 % O2 in standard incubators. hiPSC were grow on matrigel-coated dishes prepared as follows: matrigel stock solution (Corning, 354248) was diluted 1:40 in DMEM /F-12 1:1 medium (Lonza, BE12-614F and Thermo Fisher Scientific, 11765054, respectively) supplemented with P/S 100 U/mL and used to coat dishes for at least 30 minutes at 37 °C. Passages 1:6 and 1:8 were performed using ReLeSR (Stemcell technologies, 05872) or Accutase solution (Sigma-Aldrich, A6964). ReLeSR was used to detach hiPSC in clumps for expansion and standard maintenance, while Accutase solution was used for single cell passaging; in this case, ROCK inhibitor 5μM (Sigma, Y0503) was added to the culture overnight (O/N) to enhance single hiPSC survival. Cryopreservation of hiPSC was performed by single cell dissociation and storage in complete TeSR-E8 medium plus 10% DMSO supplemented with ROCK inhibitor 5μM.

The H9 human embryonic stem cell (hESC) derived neural stem cells (hNSCs) were purchased from Invitrogen (N7800-100). The hNSCs were cultured on Geltrex-coated dishes (Geltrex LDEV-free reduced growth factor membrane matrix, Thermo Fisher Scientific, A1413202) in KnockOut DMEM/F12 (Invitrogen, 12660012) supplemented with 2 mM L-Glutamine (Thermo Fisher Scientific, 25030024), 20 ng/ml EGF (Peprotech, 315-09), 20 ng/ml FGF (Peprotech, 100-18B) and 2% of StemPro Neural Supplement (Thermo Fisher Scientific, A1050801). The media was refreshed every other day and cells were routinely passaged with Accutase (Sigma-Aldrich, A6964).

### NCSC Differentiation

hiPSC were differentiated into NCSC as described in Menendez et al.^23^. NCSC differentiation required 15-20 days and was carried out as follows: 90% confluent hiPSC were detached with Accutase solution and plated on matrigel coated dishes in TeSR-E8 medium supplemented with 5μM ROCK inhibitor at a density of ∼9.2 × 104 cells per cm^2^. The day after, NCSC differentiation medium was added and changed every day for 15-20 days. NCSC medium was composed of DMEM-F-12 1:1, 10% probumin Life Science Grade from the 20% stock solution (20% m/v in DMEM F-12 1:1, stock solution; Millipore, 821001), P/S 100 U/mL, 2 mM L-Glutamine, 1% NEAA, 0.1% 1000X trace elements complex (CA055-010, Gentaur Italy Srl), 0.2% 50 mM b-mercaptoethanol, 10 μg/ml Transferrin, bovine (Holo form) (Life Technologies, 11107-018), 50 μg/ml (+)-Sodium L-ascorbate (Sigma, A4034), 10 ng/ml Heregulin-1 (Peprotech, 100-03), 200 ng/ml LONGÒR3 IGF-I (Sigma, 85580C), 8 ng/ml FGF2, 3 μM GSK3 inhibitor IX (BIO) (MedChem express, HY-10580) and 20 μm SB431542 (MedChem express, HY-10431). Cells were passaged every 4-5 days and plated at high concentration (1:1 the first time and 1:2 the following ones) on Matrigel coated dishes for the entire duration of the differentiation. Upon differentiation, NCSC were stocked as stable lines and cultured in the medium and splitting ratios. FACS following staining for HNK1 (Sigma, c6680) and NGFR (Advanced Targeting System, AB-N07) was performed to assess NCSC identity.

### RNA extraction and library preparation for RNA-seq

Total RNA was extracted from fresh pellets of iPSC or NCS using the RNeasy Mini Kit (Qiagen, 74104). Purified RNA was quantified using a NanoDrop spectrophotometer and RNA quality was checked with an Agilent 2100 Bioanalyzer using the RNA nano kit (Agilent, 5067-1512). Library preparation for RNA sequencing was performed according to TruSeq Total RNA sample preparation protocol (Illumina, RS-122-2202), starting from 250 ng - 1 μg of total RNA. cDNA library quality was assessed on Agilent 2100 Bioanalyzer, using the high sensitivity DNA kit (Agilent 5067-4626). Libraries were sequenced with the Illumina Novaseq 6000 machine at a read length of 50 bp paired-end and a coverage of 35 million reads per sample. For hNSCs, RNA was isolated using the GenElute™ Mammalian Total RNA Miniprep Kit (Sigma-Aldrich, RTN350-1KT). Total RNA was prepared using the Illumina TruSEq Stranded mRNA library Prep kit and sequenced according to the Illumina TruSeq Rapid v2 protocol on an HiSeq2500 sequencer (Illumina) at the Erasmus MC Center for Biomics at a read length of 50 bp single-end and a minimum coverage of 20 million reads per sample.

### RNA-seq analysis

Bulk RNA-seq was quantified using Salmon v 1.4 on Gencode GRCh38.p13 human genome assembly. Differential Expression Analysis was performed with edgeR library 3.38.0 using generalised linear models (GLM) using trimmed mean of M-Values (TMM) normalisation and robust estimate of the negative binomial dispersion parameter for each gene, with expression levels specified by a log-linear model, using observation weights. Master regulatory analysis was performed via recursive hypergeometric tests: for each transcription factor a gene-set derived from ChIP-seq experiments and motif-aware databases (e.g., Jaspar and TFBS) is compared to the target list of differentially expressed genes. Correction by background was performed against expressed genes. Gene Ontology (GO) and pathway enrichments were performed using topGO and clusterProfiler packages; eventual alternative background gene lists are specified in the text. Annotated heatmaps were generated using *pheatmap* library.

For hNSCs, reads of 50 bp in length were generated. Illumina adapters and poly-A-sequences were trimmed off the reads, followed by alignment to GRCh38 using HISAT2^76^. Reads were quantified using HT-seq count^77^.

### Single-cell multiomics analysis

Multiomic sequencing data was preprocessed using 10X CellRanger ARC and computationally demultiplexed based on individual-specific single nucleotide polymorphisms using SCanSNP^34^. The RNA modality was analyzed using the Scanpy package^78^ (v1.9.1). The Matplotlib (v3.4.145), and Seaborn (v0.11.146) packages were used for visualization. Pandas (v1.2.4) and Numpy (v1.22.3) were used for data handling. The ATAC modality was analyzed using ArchR suite^79^. Chromatin differential accessibility was performed with ChromVar^80^. Co-accessibility was calculated with Cicero^81^ to infer enhancer-promoter interactions and establish a dataset of regulatory regions. These regions were fed to CellOracle^44^ to calculate gene-regulatory networks, leveraging cell type-specific co-expression patterns. Master regulatory analysis was performed by measuring the enrichment of each putative TF as the proportion between the fraction of DEGs regulated over the fraction of expressed genes regulated by the same TF. Significance of each enrichment was calculated by hypergeometric test comparing the overlap between DEGs of a certain cell type and targets of a certain TFs, using expressed genes in the same cell type as background. Differential abundance was calculated on the transcriptional modality (GEX) using miloR^35^. Markers of each cell type were identified by performing cell type-specific differential expression, comparing each cell type with the rest of the transcriptome. These markers were filtered based on their cell type-specific accessibility at the promoter level, comparing each cell type against the rest of the cells in the ATAC modality. Pseudo-bulks were generated via adpbulk (github.com/noamteyssier/adpbulk), aggregating raw reads by cell type and by individual, and genes with at least 5 raw read counts in at least two individuals were filtered. Differential expression analysis was performed using previously published edgeR wrapper^53^ “edg1” for pseudo-bulk, using *estimateDisp* funciton (robust=TRUE) for dispersion estimation. FDR < 0.01 and FC >= 1.5 were used as thresholds.

### Western Blot

Cells were grown to confluence on 10-well plates, collected in PBS, pelleted by 2 min centrifugation at 400 g, and pellets were then resuspended in 100 μL RIPA buffer (50 mM Tris-HCl, pH 7.5, 150 mM NaCl, 1% Triton X-100, 0.5 mM EDTA, and 5% glycerol) supplemented with protease inhibitor cocktail (PIC), 1 mM PMSF, and 1 mM dithiothreitol (DTT). Proteins were extracted for 30 min on ice, the lysates were centrifuged at 16,000 g for 20 min at 4 °C, and the protein concentration in the supernatant was determined using the Biorad protein assay (Biorad, #5000006). For western blotting, 35 μg of protein were resolved on polyacrylamide gels, which were transferred on nitrocellulose membrane (Cytiva, 10600124), blocked for 30 min in 2.5% non-fat dry milk in TBS plus 0.05% Tween 20 (TBST), and stained with primary antibodies at 4 °C O/N. The primary antibodies used for western blotting were mouse anti-Flag (1:500, Sigma-Aldrich clone M2, F1804) and rabbit anti-tubulin (1:5,000, Abcam). Signal was detected with corresponding horseradish peroxidase (HRP)-conjugated secondary antibodies and imaged with ChemiDoc XRS+ System (Biorad).

### Digital PCR

A custom TaqMan assay was designed and ordered from ThermoFisher Scientific for each individual patient, where the ADNP reference and variant alleles were labeled as FAM and VIC, respectively. All Digital PCR experiments were performed by Cogentech SRL.

### Chromatin Immunoprecipitation (ChIP) sequencing

ChIP-seq for histone marks were performed in all iPSC lines of the cohort. Approximately 10e-7 cells were released with Accutase and then crosslinked with formaldehyde 1% in PBS 1X (Sigma, F8775) for 8 minutes at room temperature (RT) in rotation. Crosslinking was stopped by adding glycine at a final concentration of 0.125 mM and incubation on ice for 5 minutes. Cells were pelleted by centrifugation at 500 g for 3 min at 4°C and the pellet was washed once in ice cold PBS. Cells were lysed in 10 mL buffer A (50 mM HEPES pH 8.0, 140 mM NaCl, 1 mM EDTA, 10% glycerol, 0.5% NP-40, 0.25% Triton X-100) for 10 minutes on ice. After centrifugation the pellet was resuspended in 5 mL buffer B (10 mM Tris pH 8, 1 mM EDTA, 0.5 mM EGTA and 200 mM NaCl) and incubated for 5 min on ice. Then nuclei were pelleted by centrifugation at 500 g for 3 min at 4°C, resuspended in 150 µL buffer C (50 mM Tris pH 8, 5 mM EDTA, 1% SDS, 100 mM NaCl, 1x Roche complete mini protease inhibitors) and incubated on ice for 10 min. Then 3.5 mL ice cold TE buffer was added and chromatin was sheared in 13 mL tubes in a Branson Sonifier 450 (Marshall Scientific) device for 4 pulses (30 s ON / 30 s OFF) at 30% amplitude. Then 350 ul 10x ChIP buffer (0.1% SDS, 10% Triton X-100, 12mM EDTA, 167mM Tris-HCl pH 8, 1.67M NaCl) was added, chromatin was transferred into 2 mL Eppendorf tubes and spun for 10 min at 13000 g, 4°C. 1% sheared chromatin was saved as input control, the rest was transferred into fresh tubes. 5 µg of antibody were used for H3K27ac (Abcam, ab4729), H3K4me1 (Abcam, ab8895), and H3K9me3 (Abcam, ab8898) IPs while 10 µg were used for ADNP (Sigma-Aldrich, F1804 monoclonal anti-FLAG M2) and CTCF (Cell Signaling, #2899) IPs. Samples were incubated O/N on a rotating wheel at 4°C. The next morning, 40 µL Protein G Dynabeads (mixed 1:1 and washed with 1x ChIP buffer) were added and incubation was continued for another 4 h. ChIPs were washed for 1 minute each with 4x LSB (10mM Tris-HCl pH 8.0, 1mM EDTA pH 8.0, 140mM NaCl, 1% Triton X-100, 0.1% SDS, 0.1% Na-deoxycholate), 1x HSB (10mM Tris-HCl pH 8.0, 1mM EDTA pH 8.0, 360mM NaCl, 1% Triton X-100, 0.1% SDS, 0.1% Na-deoxycholate), 2x LiSB (10mM Tris-HCl pH 8.0, 1mM EDTA, pH 8.0, 250mM LiCl, 0.5% NP-40, 0.5% Na-deoxycholate), 1x TEplus (10mM Tris-HCl pH 8.0, 1mM EDTA, 50mM NaCl). Beads were transferred to a fresh tube during the last wash and wash buffer was completely removed before adding 100 µL elution buffer (10mM Tris-HCl pH 8.0, 1mM EDTA pH 8.0, 150mM NaCl, 1% SDS) and incubating 30 minutes at 65°C with constant shaking. Elution was repeated once more with 50ul elution buffer for 10 minutes and eluates were pooled, 2 ul RNaseA (20ug/ul) were added and samples were incubated for 1 h at 37°C. Input samples were adjusted to 150ul total volume with elution buffer and processed similar to ChIP samples. Then 2ul Proteinase K (20mg/ml) was added and samples were incubated 2 h at 55°C and then O/N at 65°C. 2X volumes of AMPure XP beads and 1 volume Isopropanol was added and samples were vigorously mixed and incubated for 10 min at RT. Then beads were washed twice with 80% EtOH and DNA was eluted in 30µL 10mM Tris pH8.0 for 5 min at 37°C. DNA libraries were prepared by Genomic Unit at the IFOM/IEO/IIT campus according to manufacturer protocols, and sequenced on the Illumina Novaseq 6000 instrument at 50bp paired-end read length and a coverage of 35 million reads per sample.

WT hNSCs and ADNP mutant hNSCs or F2V5-ADNP hNSCs were collected in 1x PBS with 1mM PMSF and double crosslinked with 2 mM disuccinimidyl glutarate (Thermo Fisher Scientific, 20593) by incubating for 45 min rotating at RT and with 1% buffered formaldehyde solution (50 mM HEPES-KOH, pH 7.6, 100 mM NaCl, 1 mM EDTA, 0.5 mM EGTA, 11% Formaldehyde) for 10 min rotating at RT. For single crosslinking, only 1% buffered formaldehyde solution was used. Formaldehyde was quenched by addition of 1/20 volume of 2.5M Glycine and incubation at RT for 10 min. Subsequently, the cells were washed twice with ice-cold 1x PBS with 1mM PMSF, flash frozen in liquid nitrogen and stored at −80°C. For sonication, the cell pellets were thawed on ice and resuspended in buffer A with 1x CEF (Roche, 11697498001). After 10 min of incubation under rotation at 4°C, cells were pelleted by centrifugation and resuspended in buffer B with 1x CEF. After 10 min of incubation under rotation at 4°C, cells were pelleted and resuspended in 3 ml of freshly prepared LB3 (10 mM Tris-HCl, pH 8.0, 100 mM NaCl, 1 mM EDTA, 0.5 mM EGTA, 0.1% Na-deoxycholate, 0.5% N-lauroylsarcosine) with 1x CEF and DNA was sonicated to 150-200 bp fragments using a Bioruptor Pico sonication device (Diagenode Cat# B01060001) for 14 or 10 cycles of 30 sec on and 30 sec off for double or single crosslinked chromatin, respectively. Subsequently, 1/10 volume of 10% Triton X-100 was added to the sonicated lysate which was then spun for 10 min at 1300 rpm at 4°C to remove debris. Sonicated chromatin was stored at −80°C. 10 µL of sonicated chromatin were used to check sonication efficiency and measure chromatin concentration to equalise DNA content in each condition.

For F2V5-ADNP ChIP, 200 µg of double crosslinked chromatin were diluted in ChIP-dilution buffer (1.1% Triton X-100, 0.01% SDS, 167 mM NaCl, 16.7 mM Tris-HCl pH 8.0, 1.2 mM EDTA) containing 1x CEF (Roche, 11697498001) to a final volume of 1 ml. Chromatin from WT hNSCs was used as a control. Chromatin was precleared by incubation with Protein A/G agarose beads (Santa Cruz, sc-2003) in a low binding tube for 1 hour at 4°C rotating. The pre-cleared chromatin was then transferred to a new low binding tube and incubated with 50 µL of pelleted V5-agarose beads (Sigma Aldrich, A7345) rotating at 4°C O/N. The V5-agarose beads were then washed once with Low salt wash buffer (20 mM Tris-HCl pH 8.0, 2 mM EDTA, 1% Triton X-100, 150 mM NaCl), once with High salt wash buffer (20 mM Tris-HCl pH 8.0, 2 mM EDTA, 1% Triton X-100, 500 mM NaCl), once with LiCl wash buffer (10 mM Tris-HCl pH 8.0, 1 mM EDTA, 0.25 M LiCl, 0.5% IGEPAL, 1% NaDeoxycholate) and once with T_10_E_1_ (10 mM TrisHCl pH 8.0, 1 mM EDTA). All buffers were supplemented with 1x CEF. Then, the bound chromatin was eluted from the agarose beads in ChIP elution buffer (50 mM Tris-HCl pH 7.5, 10 mM EDTA, 1% SDS) shaking at 1000 RPM for 1 hour at 65°C. The samples were de-crosslinked O/N by incubation at 55°C. DNA was isolated using Phenol/chloroform/IIA (Sigma, P3803) extraction and diluted in water.

For CTCF ChIP, 200 µg of single crosslinked chromatin with 1x CEF were incubated O/Nat 4°C rotating with either 10 µl of CTCF antibody (Merck, 07-729-25UL) or 10 µg of IgG antibody (Diagenode, C15410206) as control. Then, samples were incubated for 2 hours at 4°C rotating with 30 µl of Protein A Dynabeads (Invitrogen, 10002D) which were previously washed twice with ChIP dilution buffer. The Dynabeads were then washed thrice with Low salt wash buffer, once with High salt wash buffer, once with LiCl wash buffer and once with T_10_E_1_. The chromatin was then eluted from the Dynabeads and decrosslinked as described above for F2V5-ADNP ChIP.

For KDM1A ChIP, 200 µg of double crosslinked chromatin with 1x CEF (Roche, 11697498001) were incubated O/N at 4°C rotating with either 10 µg of LSD1 antibody (Abcam, ab17721) or 10 µg of IgG antibody (Diagenode, C15410206) as a control. For GTF2I ChIP, 100 µg of double crosslinked chromatin with 1x CEF were incubated O/N at 4°C rotating with either 5 µg of GTF2I antibody (Bethyl, A301-330A) or 5 µg of IgG antibody as a control. Histone ChIPs were performed with 30 µg of single crosslinked chromatin with 1x CEF, incubated O/N at 4°C rotating with either 1 µg of H3K4me3 antibody (Cell Signaling, 9751S) or 1 µg of IgG antibody as control. Then, samples were processed as described above for the CTCF ChIP.

DNA libraries were prepared using the ThruPLEX V2 DNA sample preparation protocol from Takara Bio at the Erasmus MC Center for Biomics at a read length of 50 bp single-end and a minimum coverage of 20 million reads per sample.

### ChIP-seq analysis

ChIP-seq reads were trimmed for library-specific adaptor contamination before being aligned to the GRCh38.p13 human genome assembly with Bowtie 2 (removing multi-mapping reads with SAMtools). We performed peak calling via MACS 2.1 using narrow settings for ADNP and CTCF (-q 0.05) and broad settings for histone marks (H3K27ac, H3K4me1 and H3K4me3). Peaks overlap were performed by means of BEDtools v2.28, and bound genes were defined as intersecting promoters (from 500bp upstream to 250bp downstream TSS), or intersecting enhancers (cell type-specific 4DGenome consortium peak sets, summits extended by 500bp upstream and downstream). Peaks in exclusion lists from ENCODE were removed. Top peak sets for ADNP and CTCF were chosen by filtering MACS2 Score and qValue: peaks with score equal or higher to median, and each qValue higher than first quartile was retained. Reference peak sets for histone marks were defined as present in at least half controls (3 out of 6 samples). We identified “lost peaks” as regions preferentially found in CTLs (i.e., regions in at least 5 out of 6 controls, and none of HVDAS, plus regions found in all controls and at most 1 out of 3 HVDAS lines), and “gained peaks” as regions preferentially found in HVDAS lines (following the same logic). Significance reported in the main text for overlaps between sets of peaks and sets of genes derived from peaks were performed via hypergeoetric test. Motif enrichments were performed with Homer v.4.11 using default parameters. Principal Component Analysis (PCA) was performed and represented using DeepTools v3.5 package on multiBamSummary output using 100bp binning. Heatmaps were generated using deepTools plotHeatmap on bamCoverage (RPGC normalization) or bamCompare (normalization on Inputs) outputs, as stated in the text. Reference peak files were chosen depending on histone marks or TF, as stated in the previous sections, or as specified in the text. Unless otherwise specified in the text, the number of mapped reads was used as the library size, mean difference genewise exact test was used for assessing the statistical significance of the intensive changes in ChIP-seq signals in each differential analysis, using edgeR 3.38.0. Master regulatory analysis was performed as it follows: for each transcription factors a gene-set derived from ChIP-seq experiments and motif-aware databases (e.g., Jaspar) is compared via hyper-geometric test to the target list of genes, correcting by background (e.g., expressed genes for differentially expressed genes in RNA-seq, or bound genes against differentially marked regions in ChIP-seq). GO and pathway enrichments were performed using ChIPseeker and clusterProfiler packages; background genes are specified in the text.

For ChIP-seq datasets from hNSCs, the reads were first trimmed by removing the Illumina adapters, followed by alignment to the reference genome GRCh38 using HISAT2^76^. After alignment, low-quality, duplicated fragments and fragments that exceed 150 bases in length were removed.

### CRISPR-Cas9-mediated tagging of ADNP in iPSCs

We designed a construct harbouring: i) hygromycin resistance gene ii) P2A self-cleaving peptide between the hygromycin resistance gene and the downstream *ADNP* open reading frame, iii) 3XFLAG tag at the 5’, iv) 5’ and 3’ 1000 bp-long homology arms (HA). This construct was electroporated along with the Cas9 and the appropriate gRNA (see supplementary material), assembled as a ribonucleoprotein (RNP) complex. The selected protospacer adjacent motif (PAM), was located in the vicinity of the *ADNP* endogenous ATG in order to enhance the homologous recombination at the starting of exon 3, where the translation of ADNP begins. Finally, because the 3’HA falls in the coding sequence (exon 3), one base of the arm that encompasses the PAM was changed by a synonymous mutation in order to avoid the cut of the construct or the newly edited region, while preserving the translational frame.

The construct was synthesised by GeneArtTM (Thermo Fisher Scientific). sgRNAs design was performed using the CRISPR design tool of Benchling available at the following website: https://www.benchling.com/. sgRNA was chosen according to two quality scores: the on-target and the off-target score, both ranging from 0 to 100, defined based on an algorithm by Doench et al^82^ and Hsu et al^83^. sgRNA was synthesised using the GeneArtTM Precision gRNA Synthesis Kit (Thermo Fisher Scientific, A29377) by *in vitro* transcription. Oligonucleotides were synthesised by Sigma-Aldrich. The concentration of the *in vitro* transcribed sgRNA was determined using a NanoDrop spectrophotometer. sgRNA/Cas9 electroporation conditions were standardised using a Cas9 purified by the Cogentech biochemistry facility (Cogentech SRL). Cas9 was produced solubilised in transduction buffer (5x Transduction buffer: 500 mM NaCl, 25 mM NaH2PO4, 250 mM NDSB-201, 150 mM glycerol, 75 mM glycine, 1.25 mM MgCl2, 1 mM 2-mercaptoethanol at pH 8.0 in milliQ water. The Neon Transfection System (Thermo Fisher Scientific) with the 100 μL kit (Thermo Fisher Scientific, MPK10096) was used to electroporate hiPSC. 70-80 % confluent cells were pretreated for 2-4 hours with 10 μM ROCK inhibitor to enhance cell-survival after electroporation. At the moment of the electroporation, hiPSC were detached using Accutase solution (1 mL x 10 cm plate), centrifuged for 3 minutes at 160g with a standard centrifuge. Cells were then resuspended in PBS and counted with an automated cell counter. The ribonucleic complex Cas9/sgRNA (molar ratio: 1:2,58) was assembled and incubated 10 minutes at 37°C to enhance the complex formation. Once formed, the complex is stable for two hours. For each electroporation reaction, 4×10^5^ cells, 1.5 μg recombination plasmid and corresponding amounts of Cas9/gRNA complex were used (10 μg of Cas9 for 4×10^5^ cells). Cells were resuspended in electroporation buffer T (provided by the kit) in order to have 4×10^5^ cells in 50 μL. The mixture was added to 50 μL of cells and then brought to 120 μL final volume adding buffer T. The tip was introduced into the electroporation station containing 3 mL of buffer E2 (provided by the kit). Electroporation was performed with the following conditions: amplitude: 900 V, pulse width: 20 ms, n° of pulses: 3. Electroporated cells were directly plated on matrigel coated dishes in TeSR-E8 supplemented with 10 μM ROCK inhibitor. Electroporated cells were plated in a 15 cm in a dish to allow clonal growth. Hygromycin B (Merck, 31282-04-9) selection was carried out after 48h from the electroporation at a concentration of 50 μg/mL for 10 days (previously tested on the same iPSC lines). After 10-15 days individual colonies were manually picked and plated in 48 well plates for expansion, screening and freezing. To extract DNA, cells were incubated O/N at 60°C with Bradley Lysis Buffer (10 mM Tris-HCl pH 7.5, 10 mM EDTA, 0,5% SDS, 10 mM NaCl in ddH2O) containing Proteinase K (1 mg/ml). The day after, ice-cold EtOH/NaCl mix (EtOH 100%, NaCl 5M) was added to precipitate the DNA and the cells were incubated 30 minutes at −80°C and then centrifuged 20 minutes at 180 g. The pellet was washed twice in cold 70% EtOH and centrifuged for 10 minutes at 180 g. DNA was eluted in 30 μL of warm TE buffer (Tris-HCl pH8, EDTA 1 mM pH 8.0 and quantified using Nanodrop spectrophotometer. 25 ng of genomic DNA were used for PCR amplification in order to screen for correctly edited clones. Further validation to ascertain the correct knock-in of the recombinant cassette was performed using Sanger sequencing. All Sanger sequencing experiments were performed by Cogentech SRL. Primers used in the screening and validation are listed in the supplementary material.

To tag ADNP in hNSCs, the gRNA (5’-ACTTCCTGTCAACAATCTTG-3’) targeting the N-terminus of ADNP was designed using the online webserver crispr.mit.edu and cloned in the pSpCas9(BB)-2A-Puro plasmid (PX459, Addgene). To integrate the F2V5-tag at the N-terminal end of the ADNP protein, we designed a 200 bp single-stranded oligodeoxynucleotide (ssODN) template containing the F2V5-tag flanked by approximately 50 bp of homology arms (see supplementary material). The hNSCs (3.5×10^6^) were transfected with 3 µg of PX459 plasmid containing the gRNA and 1 µg of the homology template by electroporation with program A33 of the Amaxa nucleofector I and the Cell Line Nucleofector™ Kit V (Lonza, catalog # VVCA-1003). 24 hours after transfection, cells were put under selection with 1 µg/ml Puromycin for 48 hours. Cells were plated at clonal density and the media was refreshed every other day until colonies were picked. The resulting candidate hNSC F2V5-ADNP clones were validated using Sanger sequencing and by checking the expression of the F2V5-tagged ADNP protein by western blot (Primers listed in supplementary material). In brief, the cells were harvested and lysed with 2x Laemmli sample buffer (0.2M DTT, 4% SDS, 1M Tris HCl pH 6.8, 20% Glycerol). After sonication, the protein samples were run on a ExpressPlus Page 4 to 12% polyacrylamide gel (GenScript, M41215). Gels were then transferred onto nitrocellulose membranes (Amersham Bioscience). The membranes were blocked in 5% Fat-free milk proteins in TBS 0,1% Tween and probed with anti-Flag antibody (Sigma-Aldrich, F3165, 1:2000) and anti-V5 antibody (Invitrogen, R96025, 1:2000) O/N at 4°C. Last, the probed blots were incubated at RT with HorseRadish Peroxidase (HRP) conjugated secondary anti-mouse (GE Healthcare, NXA931, 1:4000). The protein bands were visualised on an AI-600 digital imager (Amersham). As loading control, an anti-ACTIN antibody (Chemicon, MAB1501R, 1:4000) was used.

### CRISPR-Cas9-mediated knockout of ADNP in hNSCs

The gRNA (5’-AGAATATCCGGGGGGGATC-3’) targeting intron 4 of ADNP and the gRNA (5’-AGTTATTCAGACGGTTCAT-3’) targeting exon 5 of ADNP were designed using CRISPOR^84^, and cloned into a pSpCas9(BB)-T2A-GFP expressing plasmid (PX458, Addgene) and a eSpCas9-T2A-mCherry expressing plasmid (gift from the Barakat lab), respectively. The hNSCs (3.5×10^6^) were transfected with 3 μg or 1.5 μg of each plasmid by electroporation with program A33 of the Amaxa nucleofector I using the Cell Line Nucleofector™ Kit V (Lonza # VVCA-1003). 24 hours post transfection, GFP and mCherry expressing cells were sorted by FACS and plated at low density (1000-5000 cells) in Geltrex coated 10 cm dishes in the presence of 10 mM ROCK-inhibitor (Stem Cell Technologies, Y-27632). After approximately 20 days, single colonies were picked, expanded and genotyped using PCR and Sanger sequencing (Primers listed in supplementary material). Western blotting as described above with anti-ADNP antibody (Bethyl, A300-104A-M, 1:1000) and anti-ACTIN antibody as control confirmed the absence of ADNP in knockout clones.

### FLAG affinity purification of ADNP followed by mass spectrometry

Nuclear extracts were made of hNSC expressing endogenously F2V5-tagged ADNP and from control hNSCs according to the well-established Dignam protocol^85^. To identify the protein interaction partners of ADNP in hNSCs a FLAG-affinity purification was performed on 1.5 ml of nuclear extract and analysed by mass spectrometry, as described previously^86^. In brief, 60 ml of anti-FLAG M2 affinity agarose beads (Sigma-Aldrich, A2220) were equilibrated in buffer C-100* (20 mM HEPES pH 7.6, 0.2 mM EDTA, 1.5 mM MgCl_2_, 100 mM KCl, 20% Glycerol, 0.02% NP40 and 1x CEF protease inhibitor (Roche, 11873580001) and added to 1.5 ml of nuclear extract in no-stick microcentrifuge tubes (Alpha Laboratories, LW2410). The mixture of beads and nuclear extract was rotated at 4°C for 3 hours in the presence of 225 units of Benzonase (Merck, 70664-3). After incubation, the beads were washed 5 times with buffer C-100*. The bound proteins were then eluted of the beads at 4°C with buffer C-100* containing 0.2 mg/ml of FLAG-tripeptide (Sigma-Aldrich, F4799). TCA precipitation was performed on the elutions. The precipitated proteins were then separated on a 10% NuPAGE Novex Bis-Tris gel (Invitrogen, NP0301) and stained with the Colloidal Coomassie Blue Staining Kit (Invitrogen, LC6025). The gel lanes were cut into slices and subjected to in-gel digestion with trypsin (Promega). Nanoflow LC-MS/MS was performed on an 11 series capillary LC system (Agilent technologies) coupled to an LTQ-Orbitrap mass spectrometer (Thermo Fisher). The resulting peptide spectra from the F2V5-ADNP or the control elutions were then identified by MASCOT score using the Uniprot release 2012-11 database. The following inclusion criteria for the ADNP interaction partners were used: a MASCOT score of 50 or higher, at least 3-fold enrichment of EmPAI score in the ADNP purified sample compared to the control elution and only nuclear proteins were considered (based on the Uniprot database). In total the FLAG affinity purification was repeated 3 independent times and all interaction partners that were identified in 3 out of 3 or 2 out of 3 experiments were included in the final list of identified ADNP interactors. The BioGrid database^87^ was used to verify novel interactions.

### Nuclear/cytoplasm fractionation

iPSCs from a 10cm plate were collected and washed with 1X PBS, resuspended in 500 µL of Low Salt Buffer (100mM HEPES pH6.8, 5mM KCl, 5mM MgCl2, 0.5% NP-40 complemented with PIC), and let on ice for 15 min. Cells were centrifuged at 720xg at 4°C for 5 min. Supernatant was collected as cytoplasmic fractions. Pellet was washed twice with 200 µL of Low Salt Buffer to remove residual cytoplasmic traces. Nuclei were lysed with 500 µL of High Salt Buffer (100mM HEPES pH6.8, 5mM KCl, 5mM MgCl2, 0.5% NP-40, 250mM NaCl complemented with PIC) and incubated on ice for 30 min with occasional pipetting using p1000 and p200 or a 25G syringe. The lysate was incubated for 10 min at 37C° for Benzonase treatment (∼50 units per 200 µL lysate). Nuclei were centrifuged at 720xg for 3 min at 4°C and supernatant was collected as nuclear fraction.

### Immunoprecipitations

For KDM1A and GTF2I immunoprecipitation, 250 µL of F2V5-ADNP hNSC nuclear extract with 38 units of Benzonase (Novagen) and 1x CEF protease inhibitor (Roche) were rotated with 782 ng KDM1A antibody (Cell Signaling, 2139S) or 300 ng of GTF2I antibody (Bethyl, A301-330A) for 2 hours at 4°C in no-stick microcentrifuge tubes (Alpha Laboratories). 10 µL of ProtG Dynabeads (Invitrogen, #100-04D) equilibrated with 1 ml of C-100 buffer (20 mM Hepes pH 7.6, 0.2 mM EDTA, 1.5 mM MgCl_2_, 100 mM KCl, 20% glycerol) were blocked with 0.1 mg/ml insulin (Sigma-Aldrich) and 0.2 mg/ml chicken egg albumin (Sigma-Aldrich) in C-100 buffer for 1 hour rotating at RT and washed twice with C-100 buffer. Incubated hNSC nuclear extract with antibody was added to the blocked Dynabeads and rotated for 2 hours at 4°C in no-stick microcentrifuge tubes. Bound proteins were eluted from the beads by incubating at 95°C for 5 sanger in 28,5 μL 2x Laemmli sample buffer (0.2M DTT, 4% SDS, 1M Tris HCl pH 6.8, 20% Glycerol). As a control, an IgG (Diagenode, C15410206) immunoprecipitation was performed in the same manner. The elution samples were run on a ExpressPlus Page 4 to 12% polyacrylamide gel (GenScript, M41215). Resulting gels were transferred on nitrocellulose membranes (Amersham Bioscience). The membranes were blocked in 5% Fat-free milk proteins in TBS 0,1% Tween and probed O/N with KDM1A antibody (Cell Signaling, 2139S, 1:1000) or GTF2I antibody (Bethyl, A301-330A, 1:1000) and Flag antibody (Sigma-Aldrich, F3165, 1:2000), followed by incubation at RT with HorseRadish Peroxidase (HRP) conjugated secondary anti-Rabbit antibody (GE Healthcare, NA934V, 1:4000) or with HorseRadish Peroxidase (HRP) conjugated secondary anti-mouse (GE Healthcare, NXA931, 1:4000). The protein bands were visualised on an AI-600 digital imager (Amersham).

### Omni-ATAC

ATAC-seq was performed as reported in Corces et al^88^, an improved version of the original protocol with the main advantages of higher accuracy in calling genome-wide accessible regions, and reduced mitochondrial DNA contamination. Omni-ATAC was first thoroughly standardised to achieve the best working conditions using a Tn5 prepared and purified in the IEO crystallography unit. 50,000 iPSCs were collected and centrifuged at 500g for 5 minutes in a pre-chilled centrifuge and then briefly resuspended in ice-cold ATAC Resuspension Buffer (ATAC-RSB buffer: Tris-HCl pH 7.4 10mM, NaCl 10mM, MgCl2 3mM) supplemented with NP-40 0.1%, Tween-20 0.1% and Digitonin 0.01%. Samples were incubated on ice for 3 minutes. Lysis was washed out with ATAC-RSB supplemented with Tween-20 0.1% without NP-40 or Digitonin. Nuclei were then centrifuged for 10 minutes at 500g in a pre-chilled centrifuge. To allow transposase reaction, samples were resuspended in 50ul of ice-cold transposition mixture (TD buffer 2X, MEDS-loaded Tn5 100nM, PBS 33%, digitonin 0.01%, Tween-20 0.1%) and then incubated for 30 minutes at 37 C on agitation (1000 rpm). Tn5 was pre-loaded with pre-annealed Mosaic End double-stranded (MEDS) oligonucleotides as described in Picelli et al^89^. To clean up the transposase reaction, samples were purified with Zymo DNA Clean and Concentrator-5 kit (Zymo Research), according to manufacturer instructions. Eluted tagmented DNA was PCR amplified for 5 cycles using NEBNext Master Mix (NEB) and barcoded with Unique Dual Indexes (UDIs) which mitigate sample misassignment due to index hopping during de-multiplexing. 5 ul of the total 50 ul PCR reaction were collected for qPCR quantification using Viia7 Real-Time PCR system in order to assess the right number of additional cycles required to obtain optimal complexity during library amplification. 8 final PCR cycles (5 pre-amplification + 3 extra cycles) was established as the gold standard for iPSCs samples, using the following condition: 72°C for 5 min, 98°C for 30s, and thermocycling for 98°C for 10s, 63°C for 30s and 72°C for 1 min. DNA fragments obtained at the end of library preparation underwent a double-sided size selection to remove primer dimers and fragments larger than 1000bp. To remove DNA fragments > 1000bp, 0.5X volumes of Agencourt AMPure beads XP (Beckman Coulter) were added to the samples, then incubated for 10 minutes at RT. The supernatant, containing DNA fragments < 1000bp, was transferred in a new tube and incubated for 10 minutes at RT with 1.3X original volume AMPure beads. Supernatant, containing primer dimers, was discarded and the DNA-beads complex was washed 3X with Ethanol 80% and eluted in water. Libraries were quantified by Qubit DNA High sensitivity (Thermo Fisher), checked with Bioanalyzer high-sensitivity kit, and sequenced on an Illumina NovaSeq 6000 at 50bp paired-end read length and a coverage of 60 million reads per sample.

### ATAC-seq analysis

Reads were aligned using bowtie 2 on the same human hg38 reference used for ChIPseq. Fragment length distributions were plotted to verify the quality of each sequencing run. Reads were aggregated by genotype. Nucleosome free, mono- and di-nucleosome reads were extracted depending on their size. Reads shifting was performed using DeepTools alignmentSieve. Peak calling was performed using Genrich ATAC mode (https://github.com/jsh58/Genrich). Motif enrichment, differential peak calling between patients and controls, and quantitative analyses were performed as for ChIP-seq.

### Cortical organoids culturing, clearing and immunostaining

Cortical organoids were generated using an adaptation of the previously described protocol published by Trujillo et al^34^. Clearings and immunostainings were performed using reagents and manufacturer’s instructions provided in the MACS^®^ Clearing Kit (Miltenyi Biotec, 130-126-719). Cortical organoids of each sample were collected from shaking 6-well plates and gently transferred in 48-well plates for fixation using cut pipette tips. Organoids were washed 3 times in PBS to remove residual medium and then fixed for 20 minutes at RT with 2 ml of paraformaldehyde (4%). Organoids were washed again three times in PBS to remove fixative solution. Up to three organoids were permeabilized together in 0.5 ml of Permeabilization Solution in 1.5 ml Eppendorf tubes for 6 hours at RT under slow continuous rotation. After incubation, Permeabilization Solution was discarded and substituted with freshly prepared 1X Antibody Staining Solution (10X Antibody Staining Solution is diluted 1:10 with sterile water beforehand) complemented with primary antibodies in a final volume of 0.4 ml of 1X Antibody Staining Solution (see **Table S5**). Up to three organoids were incubated together with gentle shaking for 40 hours at 37°C. To remove unbound antibodies, Antibody Staining Solution was discarded and replaced with fresh one without antibodies, and incubated for 30 minutes at RT with slow continuous rotation 5 times. Secondary antibodies were diluted according to manufacturer’s recommendations in a final volume of 0.4 ml of 1X Antibody Staining Solution and added to the 1.5 Eppendorf tubes containing cortical organoids and then incubated for 1 hour at RT. To remove unbound antibodies, Antibody Staining Solution was discarded and replaced with fresh ones, and incubated for 30 minutes at RT with slow continuous rotation. These steps were repeated 5 times to ensure full removal of unbound antibodies. A summary of antibody details used with this protocol is reported in the supplementary material. For the embedding, agarose gel was prepared by dissolving 1.5% agarose in double-distilled water in a microwave. Organoids were transferred to the bottom of a 15 ml Falcon tubes and agarose solution was poured on top of them once it was slightly cooled down. After gel solidification (15-20 minutes), the agarose block was cut into approximately 5×5 mm pieces, each containing up to three cortical organoids, making sure that they were located in the corner of the gel block to facilitate imaging conditions. Dehydration solutions were prepared by diluting absolute ethanol in sterile water to obtain 50% and 70% ethanol solutions containing 2% Tween-20. Up to three embedded organoids were dehydrated with a series of ethanol dilutions in 15 ml Falcon tubes at RT under slow continuous rotation: 50% ethanol was incubated for 2 hours, 70% ethanol for 2 hours, and finally 100% ethanol O/N. Newly thawed 2.5 ml of Clearing Solution was added into new 15 ml Falcon tubes containing dehydrated organoids, and incubated at RT under slow continuous rotation for 3 hours. Clearing Solution was replaced with fresh one after 3 hours, and incubation was continued for additional 3 hours. Clearing Solution was discarded and substituted with Imaging Solution to proceed with images acquisition.

### Cortical organoids images acquisition and data analysis

For measuring NPCs abundance in control and mutated lines, organoids were immunostained and cleared as described above, and then visualised in a Yokogawa Spinning Disk Field Scanning Confocal System (Yokogawa CSU-W1 25µm-50µm pinhole dual disk, Nikon, Japan), equipped with motorised stage x-y-z, and a Prime BSI camera (Teledyne Photometrics, Arizona, USA), in confocal mode. Four channel (DAPI, Nestin, Pax6, and Sox2) z-stack images of the whole organoid (Z-step intervals of 5 μm) were acquired using the Nikon NIS Elements AR software (version 5.02.03) at a 10x/0.3 magnification (dry, no binning). Images were later analysed with an open-source software (FIJI-ImageJ v2.1.0, USA), in which all organoids were measured using a semi-automated macro with the criteria of analysis fully established and applicable to all images. The DAPI channel was used in all samples to normalise all area and/or count measurements (Pax6 and Sox2; pH3). All channels analysed were transformed into binary images using available methods (“Otsu” for DAPI and Sox2; “Yen” for Pax6) and converted to masks, from which the area was later measured, and calculated individually accounting for each organoid volume. For counting cells actively undergoing mitosis in the two conditions, images were processed in the Arivis Vision 4D software (version 3.5.0, Arivis, Germany), using a fully automated custom pipeline. Briefly, to facilitate the downstream 3D workflow, DAPI and pH3 channels were processed separately, and advanced image enhancement filters were applied. For volume measurements, the DAPI channel in each organoid was later segmented using the “Li” thresholder, and objects smaller than 10000 µm^3^ were filtered out. For counting the number of pH3-positive cells, the “Blob Finder” analysis operator was used to segment cells and the following parameters were applied: 1) average diameter of 6 µm; 2) 5% probability threshold; 3) 60% split sensitivity. Finally, raw data was processed, and the number of particles was normalised to DAPI volume. All data is presented as the median of *n* different organoids for all analysis.

### Single-cell Multiome (scATAC-seq + scRNA-seq)

Cortical organoids were dissociated using Dissociation Buffer composed of 0.5% BSA, 2 mM EDTA, 0.1% Accutase, 0.4 mg/ml Collagenase/Dispase (Sigma-Aldrich, 10269638001), 0.05 mg/ml DNAse I (Sigma-Aldrich, 10104159001) diluted into final volume of PBS, and filtered with 0.22 μm filters. Few homogeneous-sized organoids were incubated for approximately 40 minutes on a rotating wheel at 37°C with 1 ml of Dissociation Buffer, manually pipetting every 10 minutes. Organoids were transferred to a new Eppendorf leaving behind undissociated pieces, and then centrifuged at 300g for 5 minutes at 4°C. Cells were resuspended in 1 ml of PBS-BSA 0.04% and filtered with 40 μm Flowmi cell strainer (Bel-Art, H13680-0040) to remove the majority of debris. Cells were counted manually and then multiplexed together with other samples to obtain a final number of 1 million cells, with equal representation from each sample (e.g., 250K cells in case of 4 samples multiplexed). This step allows to pool together different genotypes and then de-multiplexed them based on their transcriptome already profiled at iPSC stage. Multiplexed cell solution was washed twice with PBS-BSA 0.04% and pellet was resuspended in 100 μL of Lysis Buffer [10mM Tris-HCl (pH 7.4), 10mM NaCl, 3mM MgCl2, 0.1% Tween-20 (Bio-Rad, 1662404), 0.1% Nonidet P40 Substitute (Sigma-Aldrich, 74385), 0.01% Digitonin (Thermo Fisher, BN2006), 1% BSA, 1 mM DTT (Sigma-Aldrich, 646563), 1U/μL RNase inhibitor (Sigma-Aldrich, 3335399001), nuclease-free water], and incubated 5 minutes on ice. Lysis Buffer was washed away three times with 1 ml of Wash Buffer [10mM Tris-HCl (pH 7.4), 10mM NaCl, 3mM MgCl2, 1% BSA, 0.1% Tween-20, 1 mM DTT, 1U/μL RNase inhibitor, nuclease-free water]. Assuming 50% nuclei loss during lysis, nuclei were resuspended in Diluted Nuclei Buffer [1X 10x Genomics Nuclei Buffer, 1 mM DTT, U/μL RNase inhibitor, nuclease-free water] according to the reference table provided by 10X protocol appendix. The volume of Nuclei Diluted Buffer is critical to fit the right range of concentration based on the number of targeted nuclei recovery, therefore avoiding overcrowding of the Chromium machine during tagmentation and GEM preparation steps. Resuspended nuclei were passed again through a Flowmi cell strainer and counted to check for the right concentration feasible for the number of targeted nuclei. 5,000 was the number of targeted nuclei recovery for each multiplexed sample in all the experiments (e.g., 20,000 nuclei were targeted in the reactions with 4 multiplexed samples). DNA libraries were prepared by Genomic Unit at the IFOM/IEO/IIT campus according to manufacturer’s protocol and sequenced on the Illumina Novaseq 6000 instrument at a coverage of 50,000 reads per nucleus.

### Single-cell Multiome analysis

Reads were aligned using cellranger-arc v.2.11. *In silico* demultiplexing has been performed on each batch of sequencing using single nucleotide variants (SNP) called from bulk RNA-seq using internal pipelines (alignment with STAR on cellranger reference, followed by GATK). Multiomic data was split into Gene Expression (RNA) and Chromatin Accessibility (ATAC) components before quality control. RNA was analysed using Scanpy 1.9.1 and ATAC was analysed using ArchR release 1.0.2. They were separately filtered according to the type of observable (counts per cell, mitochondrial genes expression/percentage of mitochondrial reads, percentage of reads in peaks, TSS enrichments, etc). The two modes were normalised independently following best practices (normalization on UMI and log-transformation, following Scanpy guidelines for RNA-seq, TF-IDF for ATAC-seq following *ArchR* pipeline). Cell type annotation was performed using cell-cycle markers for G2M and S-phase provided by the Seurat v4.0 suite and the cell cycle scoring function provided in Scanpy. Label transferring has been used to verify that cells cluster coherently in ATAC and RNA. We performed two independent dimensionality reductions (UMAP) on ATAC and RNA. Cell clustering was performed via Leiden algorithm (resolution 0.5). Integration of multiple batches was performed using Harmony. Ingestion (with scanpy.tl.ingest) on published reference data^42^ to validate cell type annotation and benchmarking. Trajectory reconstruction to assess differentiation in cortical organoids was performed in the following manner: i) after cell type annotation, diffusion pseudotime^37^ was calculated setting as root the progenitor cells expressing the highest levels of MKI67 and CDC20; PAGA was performed to reconstruct the simliarity between leiden clusters^38^ and establish the order of appearence of cell states starting from cycling cells and going to cells with higher pseudotime value; iii) Draw graphs were generated leveraging PAGA output using ForceAtlas2 algorithm (https://github.com/bhargavchippada/forceatlas2). Marker genes in both RNA- and ATAC-seq modalities were identified by first grouping reads by individual and by cell type, and summing raw read counts. To do so we used edgeR and GLM like for bulk RNA-seq, contrasting each cell type against the other ones, once for the control and once for HVDAS organoids. ChromVar 1.20.2 was applied to identify cell type-specific transcription factor activity, applying the same contrasts used for RNA. To generate the heatmap of top cell type markers we selected genes and genomic regions showing FC >= 2 and FDR <= 0.001, and represented only their intersection. Only transcription factors whose expression was cell type-specific and whose regulatory loci were accessible in the same cell type-specific fashion are reported in Fig.5A. To generate gene-regulatory networks we leveraged CellOracle^44^, which works by scanning TF motifs on the full atlas of regulatory regions that we identified by measuring co-accessibility using Cicero^81^, and harnessing cell type-specific co-expression signals to generate cell type-specific (GRN).

## Supplementary material

### Supplementary Tables

**Supplementary Table 1:** Patients and controls in iPSC cohort

**Supplementary Table 2:** Differential Expression table for iPSC and hNSC

**Supplementary Table 3:** Mass-spectrometry interactors

**Supplementary Table 4:** Cell type-specfic differential activity tables

**Supplementary Table 5:** List of gRNAs, antibodies and oligonucleotides used in this study

